# Fulfilling Koch-like postulates for animal fungal mutualists: New fungal symbionts and gallery and mycangial colonization by *Xyleborus affinis* ambrosia fungi

**DOI:** 10.64898/2026.04.23.720448

**Authors:** Abolfazl Masoudi, Esther Tirmizi, Mateo J. Valdiviezo, Ross A. Joseph, Nemat O. Keyhani

## Abstract

Fungal-animal mutualisms remain significantly understudied, yet they represent some of the most successful partnerships known in nature. Fungal farming ambrosia beetles cultivate a consortium of fungal partners that include obligate filamentous members and yeasts. These fungi are maintained in highly specialized insect organs, termed mycangia, and are cultivated as food along the beetle galleries elaborated within host trees. Here, we isolated fungi from the mycangia of *Xyleborus affinis* ambrosia beetles using both standard ethanol-wash and ethanol-free protocols. Omitting the ethanol wash significantly increased fungal recovery and diversity. We identify two previously described filamentous species, *Raffaelea arxii* and *R. fusca*, and the yeast, *Ambrosiozyma monospora*, together with two new filamentous fungi, *Neocosmospora affinis* and *Graphium ambrosium*, and two novel yeasts, *Alloascoidea xylebori* and *Wickerhamomyces ambrosius*, from both gallery walls and beetle mycangia. To meet Koch-like postulates, using mycangial colonization assays, we demonstrate that all seven fungal species were individually competent at colonizing aposymbiotic *X. affinis* mycangia, demonstrating each as viable symbionts. Our results show that ethanol-protocols can bias recovery of mycangial fungi, leading to underestimation of fungal diversity associated with ambrosia beetles. These findings provide a framework for improved characterization of ambrosia beetle-fungal mutualisms and experimental validation of fungal symbionts.

## Introduction

Ambrosia beetles are specialized wood-boring weevils (*Curculionidae: Scolytinae* and *Platypodinae*) that engage in obligate fungal farming with diverse lineages of fungi (*Ophiostomataceae*, *Ceratocystidaceae*, *Hypocreaceae*, and *Polyporales*) (ambrosia symbioses) as the sources of food for larvae and adults (Hulcr and Stelinski, 2017; Li et al., 2017; Vanderpool et al., 2018). These beetles burrow into the sapwood of trees, constructing galleries where they cultivate their fungal symbionts and rear their brood (Batra, 1966; Biedermann et al., 2013). The ambrosia beetle lifestyle is polyphyletic, with diverse unrelated lineages having adopted this fungal-animal mutualism. Due to this, different ambrosia beetle lineages have formed mutualisms with different fungal partners with varying degrees of specificity and have also evolved widely disparate means of retaining their symbionts (Kostovcik et al., 2015; O’Donnell et al., 2015; Bateman et al., 2016; Saucedo-Carabez et al., 2018; Carrillo et al., 2019; Skelton et al., 2019). Thus, the adaptation to the obligate nature of the ambrosia symbioses has led to the evolution of specialized structures on or within the beetles themselves, termed mycangia, which enable them to acquire, retain, and transport their fungal partners. The mycangial organ is responsible for sustaining mutualism between the partner microbes and the beetle host across generations and into new environments, and have arisen numerous times independently within different lineages, resulting in a wide diversity of mycangial structures in terms of shape, size, location on/within the insect body, and even distribution between males and females (Biedermann et al., 2020; Mayers et al., 2020; Spahr et al., 2020; Mayers et al., 2022). For *Xyleborus* beetle species, females (but not males), contain two preoral mycangia consisting of small pouches within the beetle head, slightly below the mandibles, and on either side of the esophagus (Hulcr and Cognato, 2010; Joseph et al., 2023; Joseph et al., 2025a). These mycangia are dynamic and contain a variety of intra-mycangial structures that may act to expand and contract the mycangia for uptake and expulsion of contents (Joseph et al., 2025a).

*Xyleborus* ambrosia beetles form mutualisms with an assortment of filamentous fungi and yeasts. These include (filamentous) species within the *Raffaelea*, *Harringtonia*, and *Dryadomyces* genera, along with members of the *Ambrosiozyma* (yeast) genus, often co-occurring or being detected during different periods of beetle/colony maturation (Campbell et al., 2016; Simmons et al., 2016; Saucedo et al., 2017). At least one filamentous partner is needed, with their nutritionally enriched conidia considered to be the main food for the beetles, although multiple fungal species are typically found in *Xyleborus* galleries, with the role(s) of any associated yeasts in particular, whether potentially as partners for the filamentous fungi and/or as food sources themselves, being less well understood (Huang et al., 2019, 2020). *X. affinis*, also known as the sugarcane shot-hole borer, is an internationally distributed ambrosia beetle found on all woodland continents, preferring humid tropical climates, and typically colonizes damp logs but can also attack trees (Sobel et al., 2015; Rodriguez-Becerra et al., 2024). These beetles show defined social behaviors, including adult and larval contributions to brood care, division of labor, with adult daughters capable of laying eggs in the natal nest and/or dispersing to sites to initiate new colonies (Biedermann et al., 2011; Biedermann, 2020). These beetles have also been used as models for examining social dynamic responses to microbial pathogens, revealing colony-level adaptations and implications of their symbiotic partners in helping combat/impede competing and/or pathogenic microbes (Grubbs et al., 2020; Masoudi et al., 2025, 2026).

The dynamic nature of the *Xyleborus* mycangia, as well as its ability to switch between (related) fungal partners, has been reported, and the use of cell cytometry has allowed for direct imaging and quantification of mycangial contents (Joseph et al., 2023; Joseph et al., 2025b). Transcriptomic responses of one partner fungus, *H. lauricola*, the causative agent of laurel wilt disease, has also been reported during fungal colonization of the mycangia, revealing temporal programs of gene expression occurring as the fungus adapts to the mycangial environment (Joseph et al., 2026). However, broader community-related dynamics remain poorly understood, and although slightly variable, one main feature of methods used to examine mycangial contents is that they include an ethanol wash, *i.e.*, of the body and/or dissected head, to minimize recovery of external “contaminating/background” microbes. Here, we used *X. affinis* as a model ambrosia beetle system to examine fungal communities associated with beetle galleries and mycangia. Specifically, we aimed to: (i) evaluate whether ethanol washing influences recovery of mycangial fungi, (ii) identify and characterize fungal taxa associated with both the galleries and mycangia of *X. affinis*, including potential new species, and (iii) confirm whether recovered fungi can colonize aposymbiotic beetle mycangia, thereby fulfilling Koch-like postulates for fungal mutualists.

## Materials and Methods

### Beetle rearing and fungal isolation from beetle galleries

Laboratory colonies of the ambrosia beetle *Xyleborus affinis* were established as described (Masoudi et al., 2025). Colony establishment was initiated by placing 2 adult females together with 1 male into sterile 50 mL polypropylene conical tubes (30 × 115 mm; FALCON^®^) containing 25 mL of an artificial sawdust-based growth medium. A total of 1 L of rearing medium was prepared using 120 g wood flour (System Three Resins, Inc., Auburn, WA, USA), 30 g coarse sweetgum sawdust (Greenville, South Carolina, USA), 40 g agar (ApeX BioResearch Products, Houston, TX, USA), 20 g sucrose (Sigma-Aldrich, St. Louis, MO, USA), 10 g corn starch (ARGO, ACH Food Companies, Inc., Memphis, TN, USA), 10 g casein (Thermo Fisher Scientific Inc., Waltham, MA, USA), 10 g yeast extract (Fisher BioReagents^TM^, Thermo Fisher Scientific, Waltham, MA, USA), 2 g Wesson salt mixture (MP Biomedicals LLC, Solon, OH, USA), and 0.7 g streptomycin sulfate (Thermo Fisher Scientific, Waltham, MA, USA). The medium was further supplemented with 10 mL of 95% ethanol and 5 mL wheat germ oil (NOW^®^ Foods, Bloomingdale, IL, USA). Colony tubes were maintained in darkness at room temperature (24°C). After 3-4 weeks of incubation, colonies were examined under a stereomicroscope to confirm gallery establishment. The microbial layer within beetle galleries was aseptically collected using a sterile toothpick and transferred into 2-mL microcentrifuge tubes containing 500 µL of sterile deionized water (dH_2_O). Sampling was conducted at two time points, 25 and 28 d, with ten galleries examined per time point. Microbial suspensions were serially diluted (1:5, 1:10, and 1:15), and each dilution was plated in duplicate onto potato dextrose agar (PDA) supplemented with 1% yeast extract (PDAY). Plates were incubated at 26°C in the dark and monitored regularly for colony development. Pure cultures were obtained using single-spore isolation. Briefly, serial dilutions of each isolate were plated onto 1% sterile water agar, and individual colonies re-streaked on PDA (filamentous fungi) or PDAY (yeasts). Fungal stocks were prepared by inoculating isolates into potato dextrose broth (PDB) for filamentous fungi or yeast-peptone-dextrose broth (YPDB) for yeasts, followed by incubation for 4 days at 25°C with shaking at 180 rpm. From these, glycerol stocks (25% glycerol, v/v) were prepared and stored at −80°C. Fungi were grown from glycerol stocks to minimize physiological changes associated with repeated subculturing on solid media. Cultures of all isolates have been deposited in the Agricultural Research Service Culture Collection (NRRL), U.S. Department of Agriculture, Peoria, Illinois, USA, under accession numbers: NRRL 65229 (*Neocosmospora affinis*), NRRL 65230 (*Graphium ambrosium*), NRRL 65231 (*Alloascoidea xylebori*), NRRL 65232 (*Wickerhamomyces ambrosius*), NRRL 65233 (*Raffaelea arxii*), NRRL 65234 (*Raffaelea fusca*), and NRRL 65235 (*Ambrosiozyma monospora*).

### Fungal isolation from beetle mycangia

Fungal communities were also isolated from the mycangia of laboratory-maintained adult beetles. Adult female beetles (28) were randomly selected from 28 different colonies. Beetles were decapitated under a stereomicroscope using sterile probes. Dissected heads were individually transferred to sterile 2 mL screw-cap tubes, and two different surface-treatment protocols were applied: (1) wash in 1 mL of 70% ethanol for 2 min, followed by three rinses in 1 mL of sterile distilled water for 1 min each, and (2) initial wash with sterile water (no ethanol), with all remaining steps identical. Aliquots from the final rinse of each protocol were plated, and the head was considered “clean” if <5 CFUs were recovered in the last wash. Following surface treatment, 200 µL of sterile water and a sterile glass bead were added to each tube. Beetle heads were homogenized using a bead beater (FastPrep-24, MP Biomedicals) at 4.0 m s^−1^ for a total of 2 min, consisting of two 1-min pulses separated by a 1-min cooling interval. The homogenate was diluted with 200 µL of sterile water, and 3×50 µL aliquots were spread onto separate PDAY plates amended with streptomycin and tetracycline (100 µg/mL each). Plates were incubated at 26°C, and the colony-forming units (CFUs) were counted. Average CFU/beetle head was calculated across triplicate plates and corrected for the dilution factor.

### Morphological observations

Fungal morphological characteristics were examined using a BZ-X1000 microscope (Keyence, Osaka, Japan) and/or an upright light microscope (Fisherbrand, Fisher Scientific, USA), the latter connected to a SeBaCam 10-MP digital camera (SeBaView software, Laxco, USA). Filamentous fungi and yeasts were cultured on PDA and PDAY, respectively. Fungal cells were transferred from actively growing cultures onto PDA/PDAY agar blocks (∼5 mm in diameter) mounted on glass slides and stained with lactophenol cotton blue. Where indicated, Z-stack images were acquired and processed using the BZ-X1000 Analyzer software to generate composite images.

### DNA extraction, PCR amplification, and sequencing

Filamentous fungal strains were cultured in PDB, and yeasts were grown in PDYA (3 d, 25°C with aeration). Cultures were centrifuged (4,000 rpm, 10 min) and washed once with sterile dH_2_O. Genomic DNA was extracted from 100 mg fungal biomass using the SPINeasy^®^ DNA Kit for Tissue with Lysing Matrix (MP Biomedicals, Irvine, CA, USA), following the manufacturer’s instructions. Multiple genetic loci were amplified for taxonomic identification. The internal transcribed spacer region of ribosomal DNA (ITS rDNA) was amplified using primers designed in this study (ITS_F and ITS_R). The β-tubulin gene (BTUB) was amplified using primers T1 and T22. A ∼1,200 bp fragment of the 3′ region of translation elongation factor 1-alpha (3′-TEF) was amplified using primers 983F and 2218R. The D1/D2 domain of the nuclear large subunit ribosomal RNA (LSU) was amplified using primers NL-1F and NL-4R, whereas the nuclear small subunit ribosomal RNA (SSU) region was amplified using primers NS-1AF and NS-8R. The two largest subunits of RNA polymerase II, RPB1 and RPB2, were also targeted for amplification. For filamentous fungi, RPB1 amplification was performed using primers DF2asc and G2R, while RPB2 was amplified with primers 5F and 7cR. For yeasts, the RPB2 region spanning exons 4-11 was amplified as two overlapping fragments, with exons 4-9 amplified using primers YRPB2-4F and YRPB2-9R and exons 6-11 amplified using primers YRPB2-6F and YRPB2-11R. The details of the primers are given in Supplementary Tables S1 and S2. All polymerase chain reaction (PCR) reactions were conducted in a total volume of 25 μL, containing 6.5 μL of 2× Taq PCR SuperMix (APExBIO, Houston, TX, USA), 2.5 μL of each primer, 2 μL of template DNA, and sterile distilled water to bring the volume to 25 μL. Amplifications were performed on a C1000 Touch thermal cycler (Bio-Rad Laboratories, Hercules, CA, USA). PCR products were visualized on 1% agarose gels, and amplicons yielding single bands of the expected size were purified using the GeneJET PCR Purification Kit (Thermo Fisher Scientific, Waltham, MA, USA). The details of the PCR conditions are given in Supplemental Table S3. Purified PCR products were sequenced using Oxford Nanopore technology (Plasmidsaurus, www.plasmidsaurus.com). Sequence reads were assembled and manually curated using BioEdit version 7.0.9.0 (Ibis Biosciences, Carlsbad, CA, USA). All generated sequences have been deposited in the National Center for Biotechnology Information (NCBI) database (Supplemental Table S4).

### Molecular identification and phylogenetic analyses of *Xyleborus* ambrosia mutualists

Preliminary taxonomic identification was conducted via BLASTn version 2.15.0 searches against the NCBI nucleotide database, which indicated that the isolates belonged to the genera *Ambrosiozyma*, *Graphium*, *Raffaelea*, *Wickerhamomyces*, *Alloascoidea*, and *Neocosmospora*, the latter characterized in our earlier work (Masoudi et al., 2025). Reference sequences for phylogenetic analyses were retrieved from NCBI based on published taxonomic frameworks (Supplemental Table S1). Sequence alignments were generated using MAFFT version 7 (Katoh and Standley, 2013) with default parameters. The most appropriate nucleotide substitution model for each alignment was selected using ModelFinder (Kalyaanamoorthy et al., 2017). Phylogenetic inference for individual loci was conducted under a maximum-likelihood framework using RAxML (Stamatakis, 2014), whereas concatenated datasets were analyzed using both RAxML and Bayesian inference under a Hidden Markov Model implemented in MrBayes (Wang et al., 2025). Analyses were performed in Topali version 2.5 (Milne et al., 2009). Maximum likelihood analyses were supported by 1,000 rapid bootstrap replicates. Bayesian analyses consisted of two independent runs of 1,000,000 generations, sampling every 1,000 generations, with the first 25% of trees discarded as burn-in. Node support was considered robust when bootstrap values ≥ 70% and posterior probabilities ≥ 0.7 were observed. Final phylogenetic trees were visualized and edited using FigTree.

### Mycangial colonization bioassays

Mycangial colonization was performed as reported (Joseph et al., 2023) with the modifications listed below. Partner fungi were grown on 100 μL of PDA in 96-well plates for 5-7 d before use. Yeast cultures were grown on PDA plus yeast extract (1%). *Harringtonia lauricola* and *Candida albicans* were used as positive and negative controls, respectively. Briefly, pupae collected from dissected colony tubes were transferred to sterile filter paper disks in Petri dishes and surface sterilized with 70% ethanol, followed by incubation for 2 minutes at room temperature, rinsing with sterile water for 1 min 3 times, and kept on sterile filter paper in a Petri dish until adult beetles emerged. Individual aposymbiotic beetles were then transferred to wells of a 96-well plate containing the test fungus. At the desired time point (24 and 48 hours), beetle heads were removed and placed individually into 2 mL tubes (with screw cap) and washed as indicated in the text (i.e., +/− 70% ethanol, followed by 3-4 water washes). Beetle heads were homogenized using a bead beater (FastPrep-24, MP Biomedicals) at 4.0 m/s for two 1-min pulses separated by a 1-min cooling interval. Microscopy and cryosectioning to visualize the beetle mycangia were performed as described (Joseph et al., 2023). Briefly, *X. affinis* beetles were embedded in Tissue-Tek optimal cutting temperature compound (OCT; Sakura) within embedding molds and rapidly cryofixed by immersion in isopentane that had been pre-cooled with liquid nitrogen. Solidified blocks were wrapped in aluminum foil, transferred to liquid nitrogen, and stored at −80 °C until sectioning. Prior to cryo-sectioning, frozen blocks were conditioned on dry ice for approximately 30 min and subsequently equilibrated in the cryostat chamber at −20 °C for 20 min. Blocks were trimmed, affixed to specimen holders, and sectioned at 16-20 μm thickness. Sections were collected directly onto glass microscope slides. Where applicable, sections were mounted using VECTASHIELD HardSet antifade mounting medium supplemented with DAPI and phalloidin (Vector Laboratories, Newark, CA, USA) to label nuclei and filamentous actin, respectively. Slides were cover slipped and cured at 4 °C for 24-48 h before imaging. Brightfield images were acquired using a BZ-X1000 microscope (Keyence Corporation, Osaka, Japan). Image stacks were reconstructed and processed in BZ-X1000 Analyzer software, and fluorescence channels were overlaid with corresponding brightfield images for visualization.

### Statistical analyses

The normality of CFU counts obtained from beetle mycangia in maintained colonies was evaluated using the Shapiro-Wilk and Anderson-Darling tests. Both raw and log_10_ (CFU + 1)-transformed data exhibited pronounced right skewness and zero inflation, deviating significantly from normality (*P* < 0.0001) (Supplemental Fig. S1). Accordingly, differences in CFU abundance between the two groups were evaluated using the non-parametric Wilcoxon rank-sum test. For CFUs obtained from mycangial colonization bioassays, normality of the raw data was assessed using the Shapiro-Wilk test. Raw CFU counts deviated strongly from a normal distribution, exhibiting pronounced right skewness and zero inflation, with Log_10_ (CFU + 1) transformation substantially improved data symmetry and reduced skewness (Supplemental Fig. S2A-B). Levene’s test indicated no significant heteroscedasticity across groups, suggesting that the assumption of homoscedasticity required for parametric analyses was not violated. All statistical analyses were conducted using JMP version 18.0.1 for Windows (SAS Institute, Cary, NC, USA), and figures were generated using GraphPad Prism version 8.0.2 for Windows (GraphPad Software, San Diego, CA, USA).

## Results

### Characterization of fungi from beetle galleries and the mycangial organ

Plating of fungi scraped from beetle gallery walls revealed a diverse collection of morphologies. Over a 6-month period, gallery wall contents were examined in 240+ colonies plated onto >1125 PDA plates, yielding 7 distinct fungal and yeast isolates, each of which were subsequently single-colony purified (Fig. 1A-H). Similarly, plating of over 120 extracted beetle heads with paired mycangia revealed the same set of seven fungal isolates, although significant variation was seen between beetles. As detailed in the Methods section, enumeration of mycangial contents involved dissection of the heads and subsequent extraction via crushing (bead beating) in buffer. Typically, to minimize carry-over or “contamination” by non-mycangial fungi, i.e., those on the surface and/or around/near the mandibles, the dissected head is subjected to an initial ethanol wash (followed by three water washes), a procedure commonly used in the literature, including our own previous work (Joseph et al., 2023). However, as our recent structural studies of the mycangia revealed that it can be “open” to the outside (Joseph et al., 2025a), we considered it likely that ethanol can penetrate into the mycangia, potentially resulting in the death/lysis of resident susceptible fungi. To test this, we performed mycangial extractions with and without initial ethanol washes. As controls for any surface and/or non-mycangial CFUs, aliquots corresponding to each wash before final extraction were plated. With or without the ethanol wash, the last water wash resulted in <1 CFU/plate (typically zero), thus the experiment was considered valid in reflecting mycangial contents. After extraction (bead beating) and plating of the washed heads, the diversity of fungal recovery was higher in the non-ethanol vs ethanol protocols (Fig. 1I).

**Figure 1.**
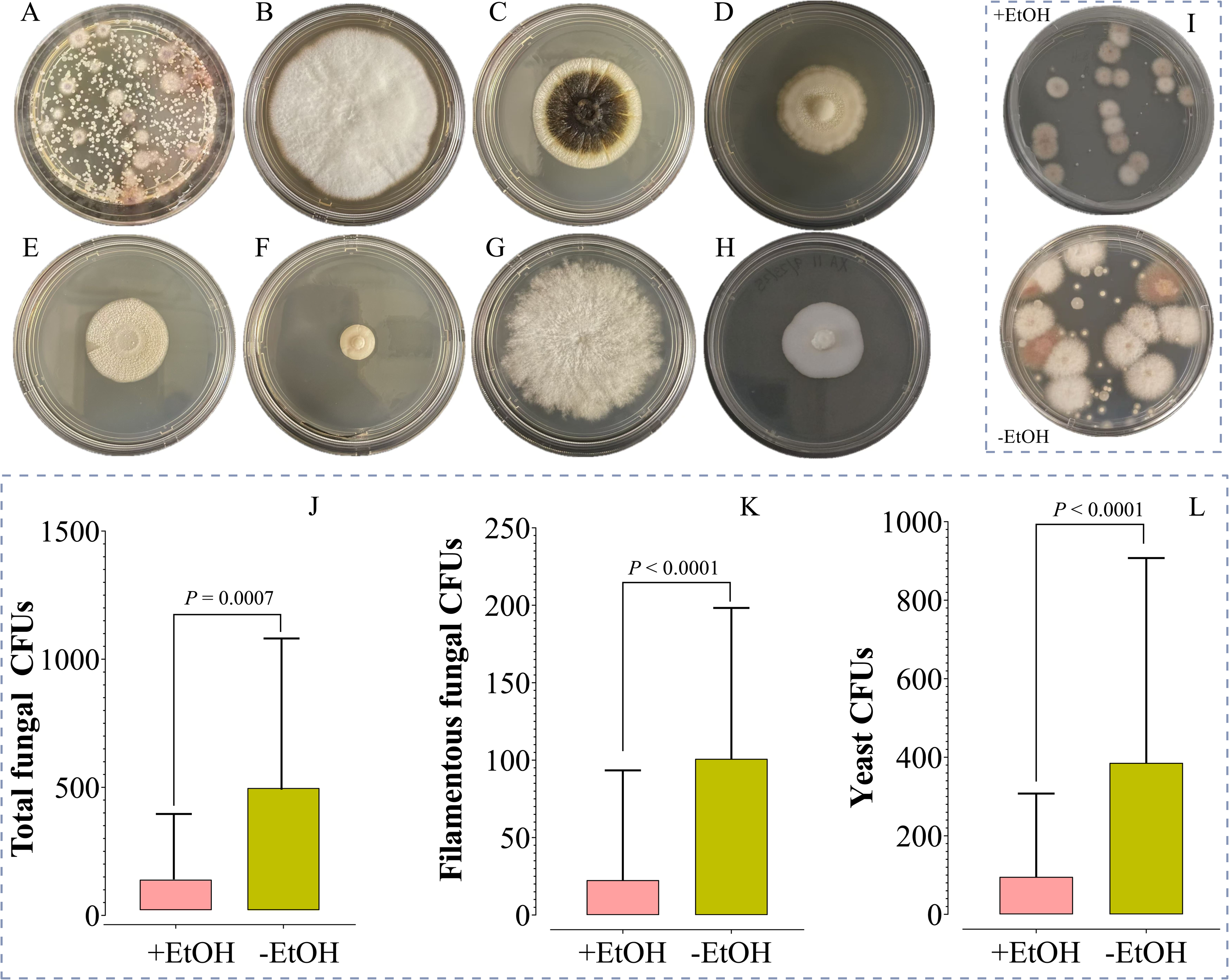
Culturable microbial communities associated with *Xyleborus affinis* galleries and mycangia. (A) Representative plate showing culturable microbial colony-forming units (CFUs) isolated from *X. affinis* galleries. (B-H) Representative colony morphologies of dominant fungal and yeast taxa isolated from *X. affinis* galleries and mycangia: (B) *Neocosmospora affinis*, (C) *Raffaelea arxii*, (D) *Raffaelea fusca*, (E) *Graphium ambrosium*, (F) *Ambrosiozyma monospora*, (G) *Alloascoidea xylebori*, and (H) *Wickerhamomyces ambrosius*. (I) Representative plates showing microbial CFUs recovered from *X. affinis* mycangia following surface treatment with ethanol (+EtOH) or without ethanol (-EtOH). (J) Total fungal CFUs recovered from mycangia under +EtOH and -EtOH treatments. (K) Filamentous fungal CFUs recovered from mycangia under +EtOH and -EtOH treatments. (L) Yeast CFUs recovered from mycangia under +EtOH and -EtOH treatments. Bars represent mean CFUs ± SD; non-parametric Kruskal-Wallis tests were used to assess statistical differences between treatments due to non-normal data distributions.

Quantification of mycangial contents revealed a significant difference in total fungal CFUs between ethanol and no-ethanol treatments (Kruskal-Wallis test, χ^2^= 11.36, df = 1, *P* = 0.0007; Fig. 1J). Beetles from the no-ethanol treatment exhibited substantially greater fungal recovery, with a median total fungal abundance of 305 CFUs (Q1-Q3: 0-647.5) compared with 0 CFUs (Q1-Q3: 0-30) in the ethanol treatment. Separation of the fungal community into filamentous and yeast-like fractions revealed similar patterns; recovery of filamentous fungi was significantly lower in the ethanol treatment than in the no-ethanol treatment (Kruskal-Wallis test χ^2^= 18.33, df = 1, *P* < 0.0001; Fig. 1K), with median filamentous fungal abundances of 0 CFUs (Q1-Q3: 0-0) and 85 CFUs (Q1-Q3: 0-177.5), respectively. Likewise, recovery of yeast-like fungi was significantly reduced in the ethanol treatment relative to the no-ethanol treatment (Kruskal-Wallis test, χ^2^ = 17.38, df = 1, *P* < 0.0001; Fig. 1L), with median yeast abundances of 0 CFUs (Q1-Q3: 0-17.5) and 130 CFUs (Q1-Q3: 30-577.5), respectively.

### Identification of seven fungal and yeast taxa isolated from beetle gallery walls and mycangia

To determine the identities of fungal isolates, the nucleotide sequences of several discriminating loci, including sections of the internal transcribed spacer (ITS), β-tubulin (*BTUB*), translation elongation factor-α (*TEF*), RNA polymerase subunit-1 (*RBP*1), RNA polymerase subunit-2 (*RBP*2), small subunit ribosomal rRNA (SSU), and/or large subunit ribosomal rRNA (*LSU*) were determined and analyzed as detailed in the Methods section. LSU sequences from one isolate clustered with *Raffaelea arxii* and from another to *R. fusca*, with strong bootstrap support in maximum-likelihood analyses, consistent with the observed >99% sequence identity to reference strains (Supplemental Fig. S3). Similarly, the 3′-*TEF* sequence from one yeast isolate showed > 99% sequence identity to *Ambrosiozyma monospora* (Supplemental Fig. S4). The remaining four isolates represent new species as characterized below.

#### Material examined and habitat

All four isolates described below were isolated from *X. affinis* laboratory colony gallery walls and from beetle mycangia.

***Neocosmospora affinis*** (Ascomycetes, Hypocreales) Masoudi & Keyhani, sp. nov. (Fig. 2; Supplemental Figs. S5-S9). Mycobank No.: MB861818

**Figure 2.**
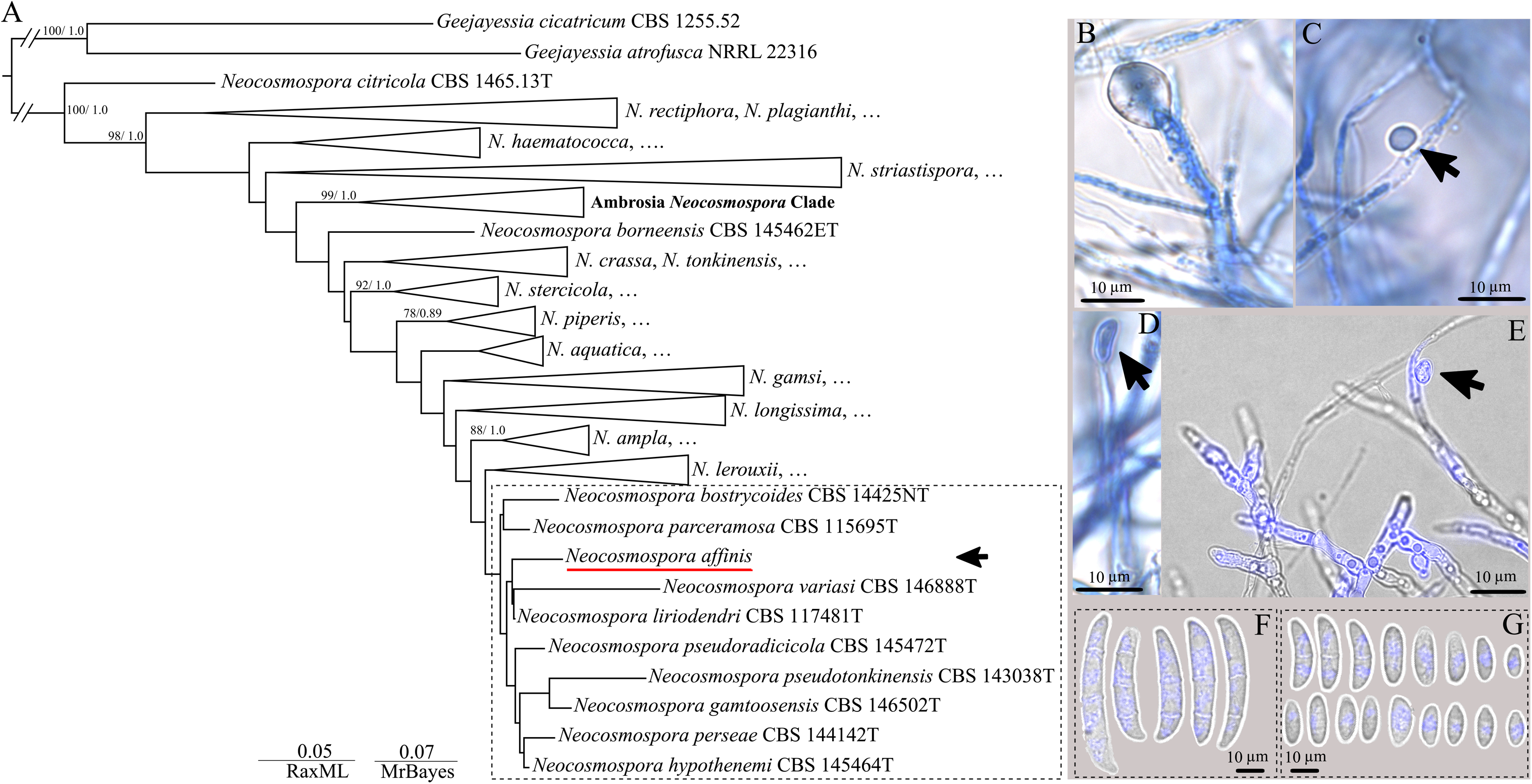
Phylogenetic placement and morphological characteristics of *Neocosmospora affinis*. (A) Concatenated phylogenetic tree inferred from combined *ITS*, *LSU*, *RPB1*, *RPB2*, and *TEF* gene sequences, showing the placement of *N. affinis* within the *Neocosmospora* clade. Support values (≥ 70% bootstrap support for maximum likelihood; ≥ 0.7 posterior probability for Bayesian inference) are indicated at branches. *Geejayessia cicatricum* and *G. atrofusca* were used as the outgroup. The dashed box highlights the phylogenetic region containing *N. affinis* and its closest related taxa (*N. variasi*, *N. liriodendri*, and related species) for easier visualization and does not represent an expansion of the Ambrosia Neocosmospora clade. The scale bar represents substitutions per site. (B-E) Chlamydospores of *N. affinis* formed on hyphae, illustrating variation in shape and position (arrows). (F) Macroconidia of *N. affinis*, showing typical fusiform shape and septation. (G) Microconidia of *N. affinis*. Scale bars = 10 µm. Note: macroscopic colony morphology for this isolate is given in Fig. 1B.

#### Etymology

“*affinis*” from beetle host *Xyleborus affinis*.

#### Phylogenetics

Phylogenetically distinct from closely related species such as *N. variasi* (MB834439) and *N. liriodendra* (MB831185).

#### Diagnosis

*N. affinis* differs from *N. liriodendri* and *N. variasi* by producing mostly aseptate to 1-septate obovoid to subcylindrical conidia on simple to sparingly branched conidiophores, with fewer and less variable chlamydospores.

#### Description

Asexual morphs: *N. affinis* exhibits a combination of morphological traits intermediate between those of *N. liriodendri* and *N. variasi*. Similar to *N. liriodendri*, conidiation in *N. affinis* is dominated by simple to sparingly branched aerial conidiophores bearing terminal monophialides, with conidia produced in false heads. Phialides are elongate and subcylindrical, and conidia are predominantly hyaline, smooth-walled, and obovoid to subcylindrical, mostly aseptate to occasionally 1-septate, lacking the pronounced falcate, multiseptate sporodochial conidia characteristic of *N. variasi*. In contrast to *N. liriodendri*, however, *N. affinis* shows greater variability in conidial size and shape, with occasional short-clavate forms and more frequent septation. Chlamydospores are present and formed terminally or intercalarily but are less abundant and less variable than those reported for *N. variasi* and lack the pronounced clustering typical of that species. Chlamydospores globose to subglobose, terminal or intercalary, solitary or in short chains, hyaline, smooth- to slightly thick-walled, approximately 5-9 µm diam. Macroconidia multiseptate, hyaline, smooth-walled, cylindrical to slightly clavate, straight to gently curved, with rounded to slightly tapered ends, approximately 30-45 × 5-7 µm. Microconidia aseptate to occasionally 1-septate, hyaline, smooth-walled, cylindrical to subcylindrical, straight, 6-10 × 2-3 µm, formed abundantly. Sexual morph not observed.

#### Culture characteristics

On PDA, colonies of *N. affinis* are white to pale cream with dense aerial mycelium and a regular margin. Colonies on PDA-covered 90 mm plates after 5 days at 25°C, with a growth rate of 13.8-14.1 mm/day.

***Graphium ambrosium*** (Ascomycetes, Microascales) Masoudi & Keyhani, sp. nov. (Fig. 3; Supplemental Figs. S10- S14). Mycobank No.: MB861819

**Figure 3.**
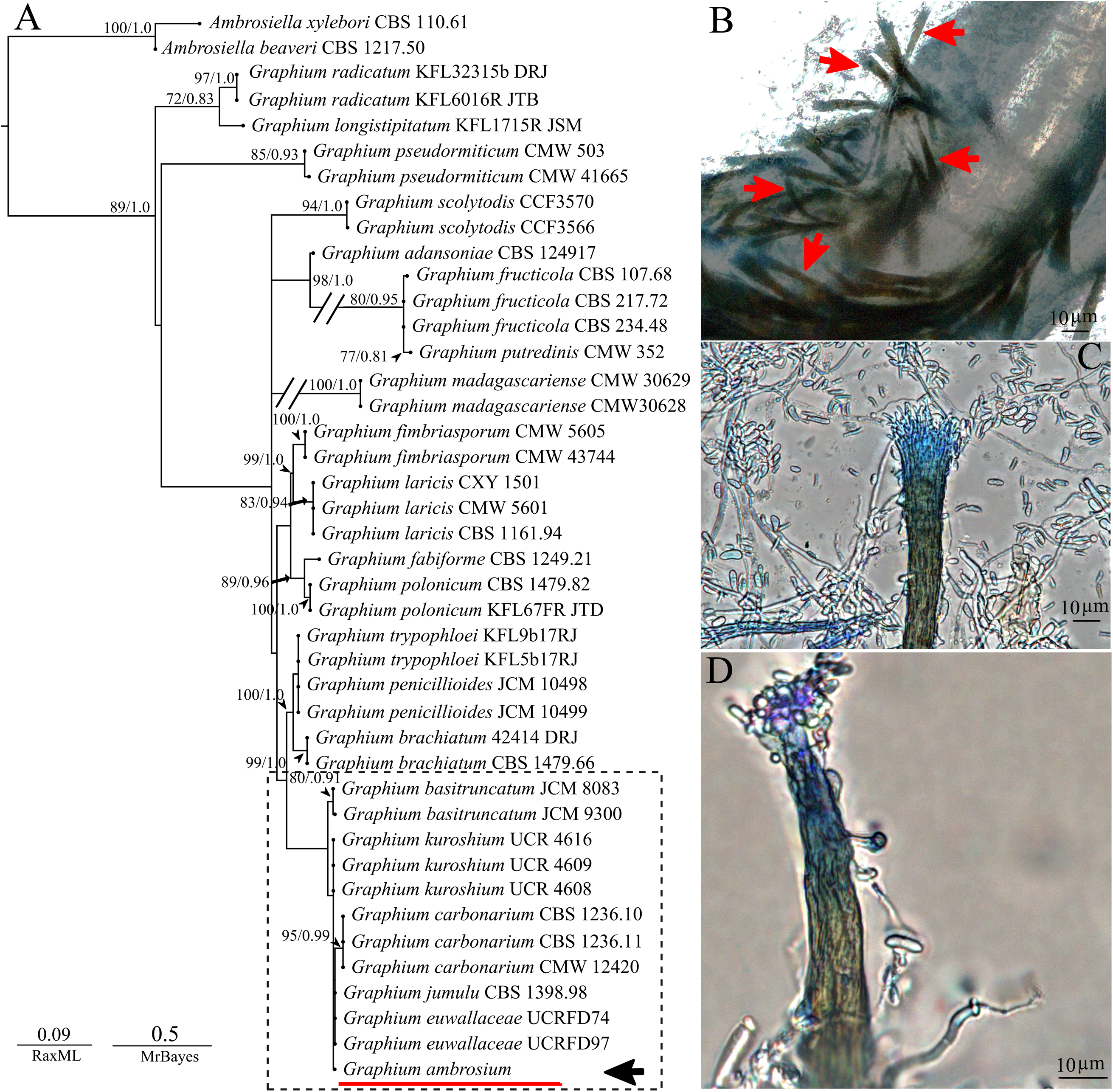
Phylogenetic placement and morphological characteristics of *Graphium ambrosium*. (A) Concatenated phylogenetic tree inferred from combined *SSU*, β*-tubulin* (*BTUB*), *LSU*, *ITS*, and *TEF* gene sequences, showing the placement of *G. ambrosium* within the genus *Graphium*. Support values (≥ 70% bootstrap support for maximum likelihood; ≥ 0.7 posterior probability for Bayesian inference) are indicated at branches. The *G. ambrosium* isolate is highlighted. The scale bar represents substitutions per site. (B-D) Morphological characteristics of *G. ambrosium* cultured on potato dextrose agar (PDA), showing elongated, slender synnemata bearing obovoid to cylindrical conidia (arrows). Scale bars = 10 µm. Note: macroscopic colony morphology for this isolate is given in Fig. 1E.

#### Etymology

“*ambrosium*” from the ambrosia beetle host.

Phylogenetics: *Graphium ambrosium* is placed within the species complex of the *Graphium* clade and groups in a well-supported clade together with *G. basitruncatum*, *G. kuroshium*, *G. carbonarium*, *G. jumulu*, and *G. euwallaceae* (dashed box in Fig. 3). Within this clade, *G. ambrosium* forms a distinct lineage, supporting its recognition as a novel species.

#### Description

Asexual morphs: Sexual morph not observed. Asexual structures are characterized by the formation of synnemata. Synnemata abundant, arising singly or in small groups, erect, darkly pigmented, composed of compactly aggregated conidiophores, with the apex expanding into a conidiogenous head (Fig. 3B-D). The conidiogenous apparatus is well developed at the synnematal apex, bearing numerous conidiogenous cells. Conidiogenous cells annellidic, producing hyaline conidia. Conidia aseptate, smooth-walled, hyaline, cylindrical to slightly clavate, accumulating around the synnematal apex (Fig. 3C, D). Sexual morph not observed.

#### Culture characteristics

On PDA, colonies started out hyaline and gradually developed an ivory-white to light yellow coloration as they matured. Growth was circular overall, with gently uneven margins. The colony surface appeared dry and compact, with hyphae closely appressed to and mostly embedded within the agar. Growth was optimal at 25 °C, with an average radial expansion rate of 8 ± 0.3 mm/day.

***Alloascoidea xylebori*** (Ascomycetes, Saccharomycetales) Masoudi & Keyhani, sp. nov. (Fig. 4; Supplemental Figs. S15-S19). Mycobank No.: MB861820

**Figure 4.**
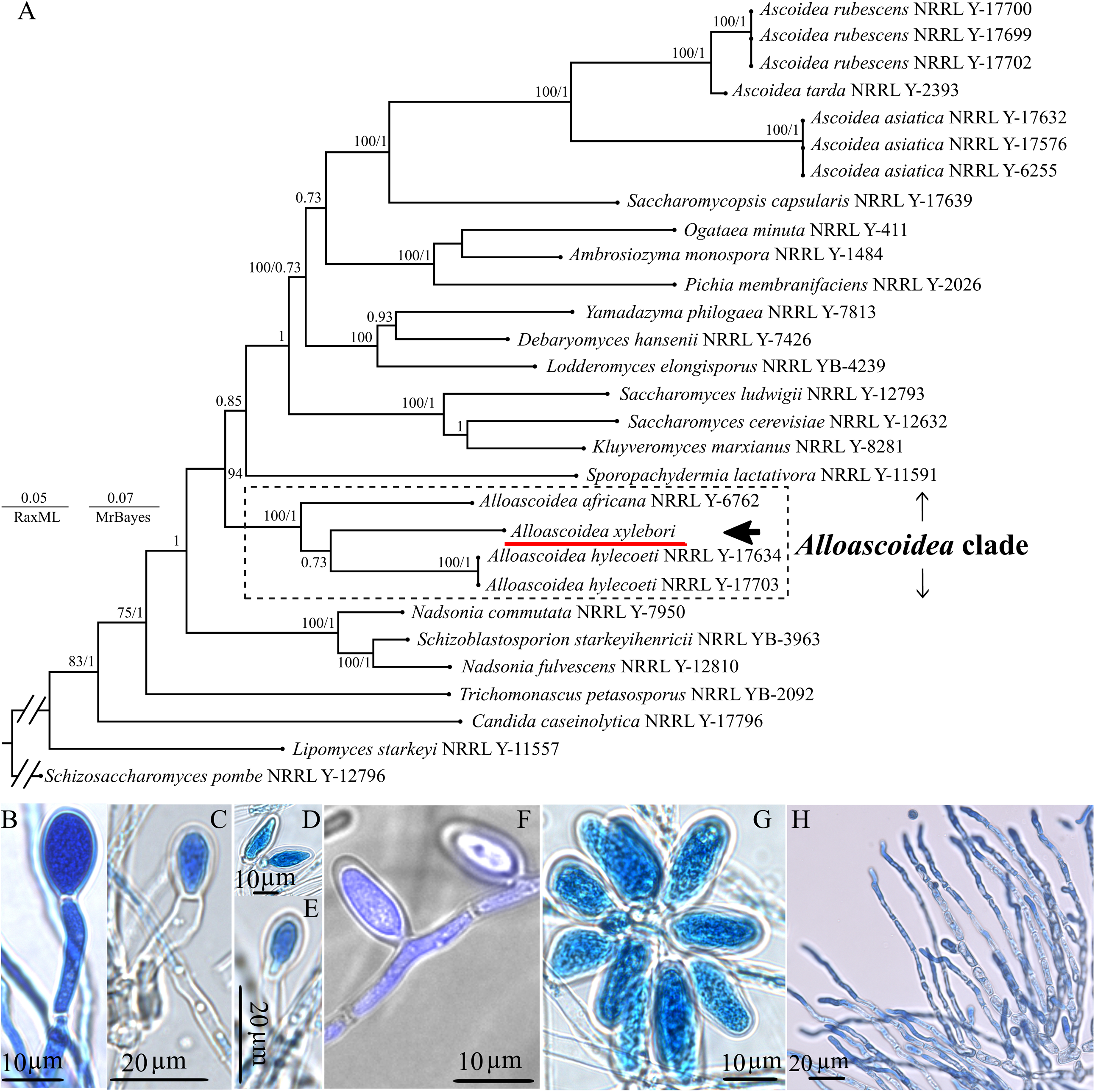
Phylogenetic placement and morphological characteristics of *Alloascoidea xylebori*. (A) Concatenated maximum likelihood and Bayesian phylogenetic tree inferred from *LSU*, *SSU*, *TEF*, *RPB1*, and *RPB2* gene sequences. Numbers at nodes indicate bootstrap support values (RaxML) and posterior probabilities (MrBayes). The *Alloascoidea* clade is highlighted, with *A. xylebori* indicated by an arrow. *Schizosaccharomyces pombe* was used as the outgroup. Scale bars represent substitutions per site. (B-F) Light micrographs showing asci of *A. xylebori* at different developmental stages, illustrating ascus shape and ascospore arrangement. (G) Cluster of mature asci emerging from ascosporogenous hyphae. (H) Expansion and elongation of hyphae into long ascosporogenous hyphae. Scale bars are indicated in each panel. Note: macroscopic colony morphology for this isolate is given in Fig. 1G.

#### Etymology

“xylebori” from beetle host *Xyleborus affinis*.

Phylogenetics: In the concatenated phylogenetic analysis, *A. xylebori* clustered within a well-supported *Alloascoidea* clade together with *A. africana* (MB802506) and *A. hylecoeti* (MB802504), confirming its placement within the genus *Alloascoidea* (Fig. 4A).

#### Diagnosis

This distinguishes *A. xylebori* from *A. africana*, where budding growth is prominent. Description: Sexual morph observed. Asci are produced from elongated ascosporogenous hyphae. Early developmental stages show solitary or paired asci arising terminally from hyphal tips (Fig. 4B-F). During development, hyphae expand and elongate into conspicuous ascosporogenous hyphae (Fig. 4H), from which clusters of mature asci emerge (Fig. 4G). Asci hyaline, smooth-walled, elongate to clavate, approximately 12-20 µm long and 4-7 µm wide, containing multiple ascospores arranged longitudinally. At maturity, asci rupture or deliquesce apically, releasing ellipsoidal to ovoid, hyaline ascospores. Budding cells were not observed under the conditions examined.

#### Culture characteristics

On PDA+Y (1% w/v), colonies grow rapidly, producing dense, cottony to felty mycelium. At 25 °C, colonies expand at an average rate of 17.0 ± 0.4 mm/ d, reaching the edge of a 90-mm Petri dish within 5-6 days. Colony surface remains pale cream to off-white, with abundant aerial hyphae developing with age.

***Wickerhamomyces ambrosius*** (Ascomycetes, Phaffomycetales) Masoudi & Keyhani, sp. nov. (Fig. 5; Supplemental Figs. S20-S24). Mycobank No.: MB861835

**Figure 5.**
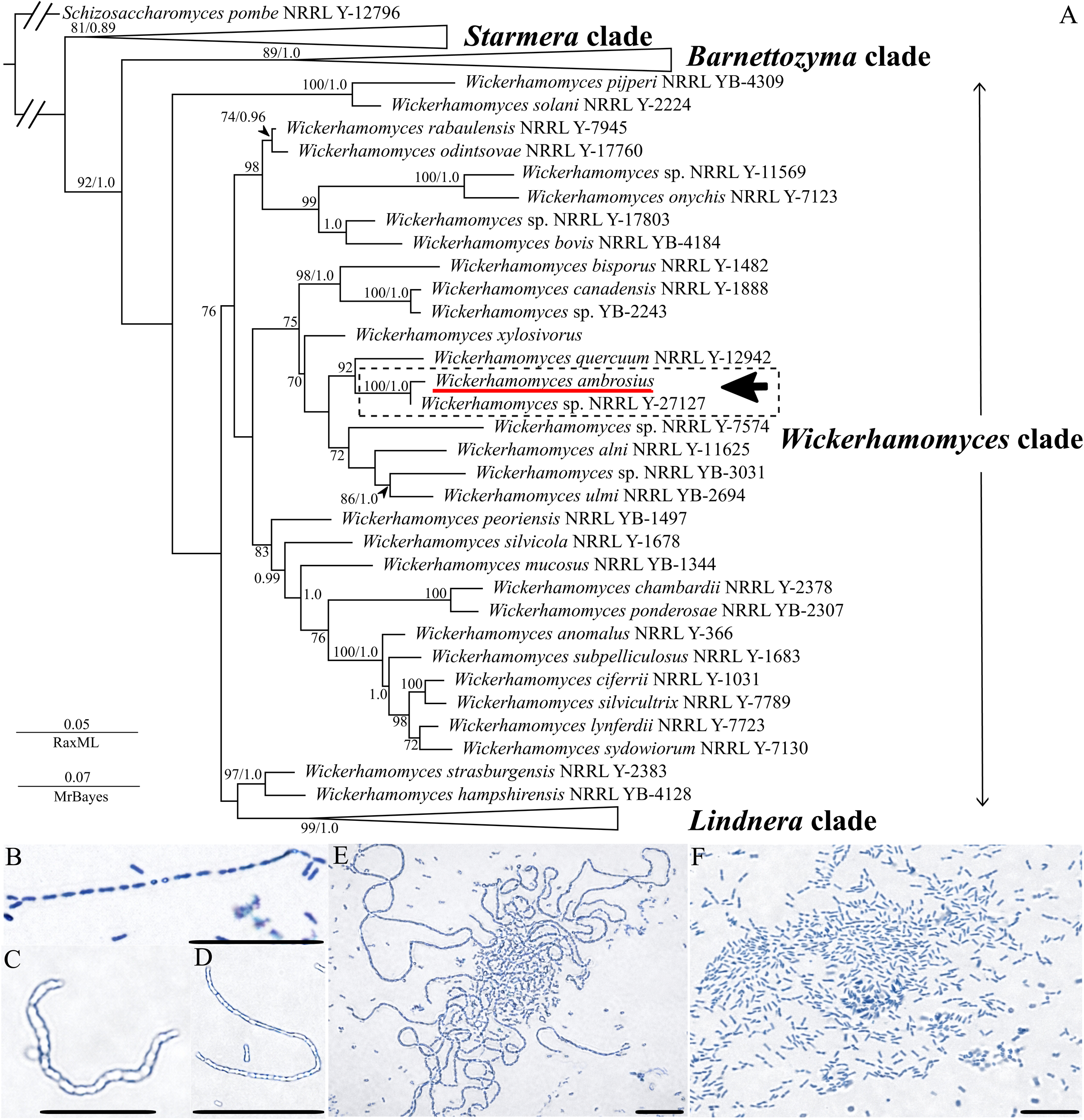
Phylogenetic placement of *Wickerhamomyces ambrosius*. (A) A concatenated maximum likelihood and Bayesian phylogenetic tree was inferred from *ITS*, *LSU*, *SSU*, and *TEF* gene sequences. Numbers at nodes indicate bootstrap support values (RaxML) and posterior probabilities (MrBayes), respectively. The *Wickerhamomyces* clade is indicated, with *W. ambrosius* highlighted. Major related clades (e.g., *Barnettozyma*, *Lindnera*, and *Starmera*) are labeled for reference. *Schizosaccharomyces pombe* was used as the outgroup. The scale bar represents substitutions per site. (B) Chains of budding yeast cells showing multilateral budding. (C-D) Elongated yeast cells and pseudo-hyphal-like elements were observed under light microscopy. (E) Aggregated pseudo-hyphal growth forms intertwined filamentous structures. (F) Abundant single-celled yeast morphology with oval to elongate cells. Scale bars = 15 μm (B-F). Note: macroscopic colony morphology for this isolate is given in Fig. 1H.

#### Etymology

“ambrosius” from ambrosia beetle, *Xyleborus affinis*.

#### Description

Multilateral budding on a narrow base. The cells are spherical, ovoid, or elongate in shape. Pseudohyphae and long growing chains were observed. Asci are persistent or deliquescent and form one to four ascospores, which may be hat-shaped or spherical and bear an equatorial ledge.

Culture characteristics: On PDA+Y (1% w/v), colonies developed slowly compared with the other isolates described in this study. Colonies were pale cream to off-white, compact with a raised central region, and exhibited a smooth surface with regular margins. Growth remained comparatively limited throughout the observation period. At 25 °C, colonies expanded at an average growth rate of [0.3 ± 0.01] mm/day.

### Fulfilling Koch-like postulates

To confirm that the isolated fungi are indeed mycangial mutualists (i.e., capable of colonizing the mycangia), all seven identified fungal taxa, as well as the previously characterized mycangial mutualist, *H. lauricola*, and the non-ambrosia fungus, *C. albicans*, were fed to aposymbiotic *X. affinis* beetles as detailed in the Methods section. Two mycangial colonization time points (24 and 48 h) and beetle head extractions with and without an ethanol wash step before maceration were performed (Fig. 6). An aliquot of the last water wash of the dissected heads was also plated to confirm minimal non-mycangial CFUs, and in all cases, < 1 CFU/head was detected. The CFU data showed an unnormalized distribution (Supplemental Fig. S25) and, to allow for statistical analyses, were log10-transformed (CFU + 1). These data indicated that all eight fungal symbionts, but not the non-symbiont, *C. albicans*, could colonize the mycangia of *X. affinis*, albeit with significant variation in response to ethanol, time, and levels (CFUs) recovered. For both yeasts and filamentous fungi, an ethanol wash significantly reduced recovery, with some taxa more sensitive than others.

**Figure 6.**
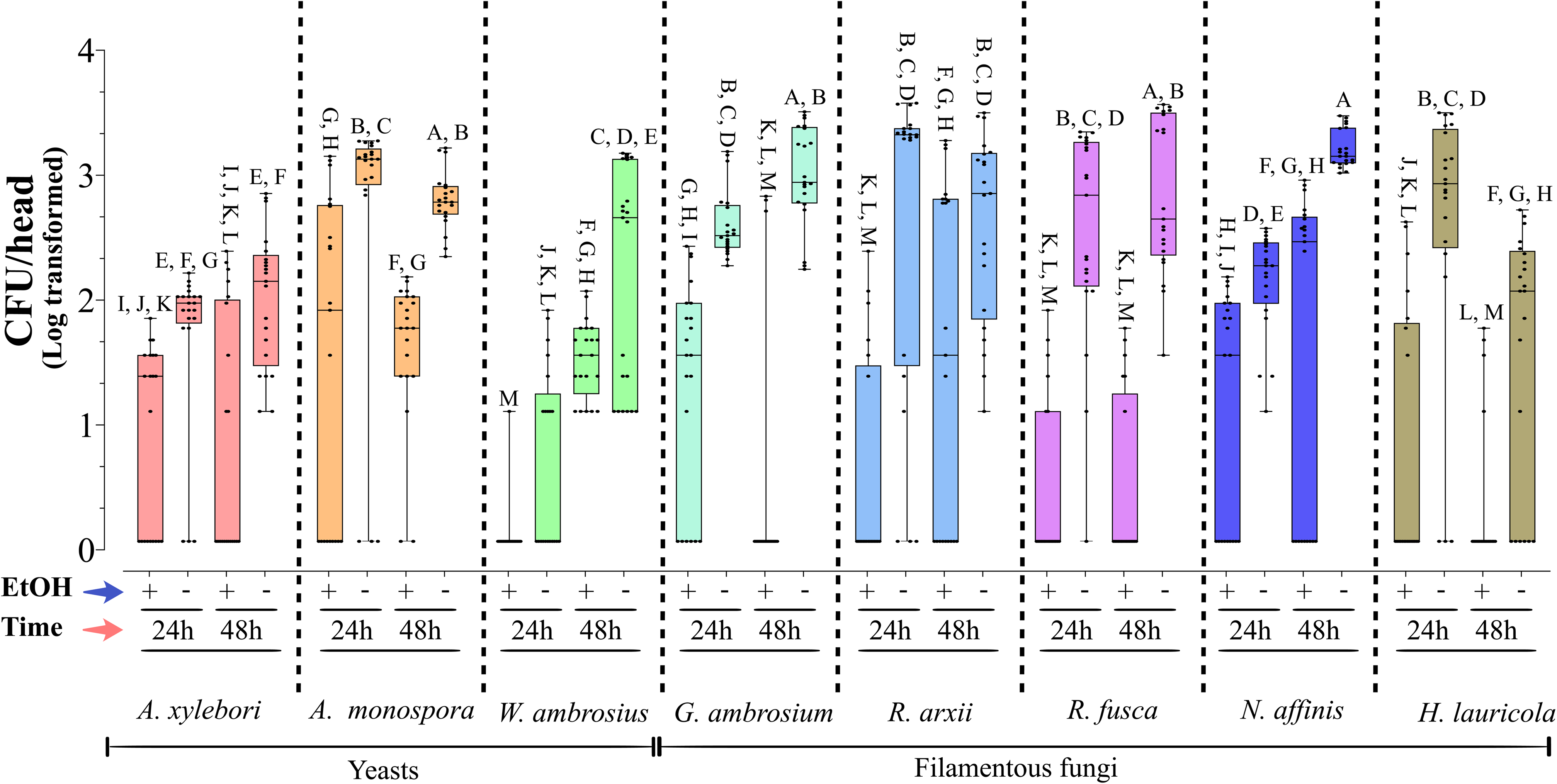
Mycangial colonization by yeast and filamentous fungal isolates in aposymbiotic ambrosia beetles. Mycangial colonization bioassays were conducted by individually exposing aposymbiotic adult beetles to PDA wells pre-inoculated with test fungi for 24 or 48 h. Following exposure, beetle heads were dissected, surface-treated with or without ethanol (EtOH), homogenized, and aliquots plated to quantify fungal recovery. Colony-forming units (CFUs) recovered per head were log-transformed and normalized prior to analysis. Seven fungal isolates were evaluated, including the yeasts, *Alloascoidea xylebori*, *Ambrosiozyma monospora*, *Wickerhamomyces ambrosius*, and the filamentous fungi, *Graphium ambrosium*, *Raffaelea arxii*, *Raffaelea fusca*, and *Neocosmospora affinis*. *Harringtonia lauricola* was included as a positive control due to its established ability to colonize beetle mycangia. Boxplots represent the distribution of normalized log_10_ (CFU/head) values, with individual data points shown, and include the median (center line), interquartile range (box), and data range (whiskers). Because datasets include zero-inflated and right-skewed values, the median may differ from the arithmetic mean, and the median should not be interpreted as a mean value. Different letters above boxes indicate statistically significant differences among treatments and time points (*P* < 0.05; ANOVA followed by Tukey-Kramer post hoc tests).

The yeast *A. xylebori* was highly sensitive to ethanol, with moderate but stable levels of colonization seen over time: = 0.824 ± 0.16 (SE) and 0.77 ± 0.21 (SE) CFUs/head at 24 and 48 h, respectively. *A. monospora* showed a significant difference +/− ethanol, although even with the ethanol wash, robust numbers of colonies were recovered. Similar to *A. xylebori*, the 24 and 48 h time points showed similar levels of mycangial colonization (+/− ethanol, but significantly higher recovery with no ethanol wash). *A. xylebori* proliferation within the mycangia was significantly higher than *A. monospora*, and along with *G. ambrosium* and the two *Raffaelea* species, showed the highest across both time points mycangial colonization (CFU counts). At 24 h, almost no *W. ambrosius* could be detected in ethanol-washed samples, with low overall counts of 0.049 ± 0.22 (SE) CFUs/head even in the absence of an ethanol wash. By 48 h, however, colonization appeared to have stabilized, although as noted, a clear ethanol effect was noted on recovery: 1.466 ± 0.066 (SE) and 2.128 ± 0.194 (SE) CFUs/head +/− ethanol wash, respectively. Both *G. ambrosium* and *R. fusca*, were very sensitive to ethanol, with little or none recovered at either 24 or 48 h colonization time points. However, omitting the ethanol wash resulted in robust, stable recovery of both fungal species from mycangia. *R. arxii* and *N. affinis* were more resistant to the ethanol wash than the other fungi tested, as was *H. lauricola* (particularly at the 24 h time point), but again, as with the other fungi tested, higher CFU recovery was clearly seen in the absence of an ethanol wash. In addition, like *R. fusca* and the yeasts, *R. arxii* recovery from mycangia was essentially equivalent at both time points; however, for *N. affinis*, the 48 h showed significantly higher CFU recovery than the 24 h time points, whereas for *H. lauricola*, the reverse was seen, i.e., higher recovery from the 24 vs 48 h time point.

To confirm colonization, sections of colonized beetles were examined for the presence of the fungal taxa within the mycangia (Figure 7). Fungal cells from all seven filamentous and yeast isolates were observed, with all producing yeast-like/blastospores within the mycangia, i.e., non-filamentous growth. Cell shapes varied, but in large part coincided in size with *in vitro* single cells; *A. monospora*, mostly uniform ∼2-3 μm diameter circular cells, *N. affinis*, a range from 2-5 μm circular cells*, G. ambrosium*, cylindrical ∼2 x 5 μm cells*, R. arxii*, circular and teardrop shaped, ∼3-4 μm cells, *R. fusca*, a range from 2-5 μm mostly circular cells, *A. xylebori*, a mixture of large ∼11-12 μm and smaller ∼3-4 μm circular/teardrop cells, and *W. ambrosius*, very small and sparsely distributed (0.5-1.0 μm) cells. Visual images of the mycangial contents when fed each fungal taxa correlated with the quantitative CFU data above, with *G. ambrosium* colonization appearing the most robust. Mycangial structures, e.g., the entry/exit channel, Ross and eyelash projections (RP and EP, respectively, (Joseph et al., 2023; Joseph et al., 2025a)) were noted in many images.

**Figure 7.**
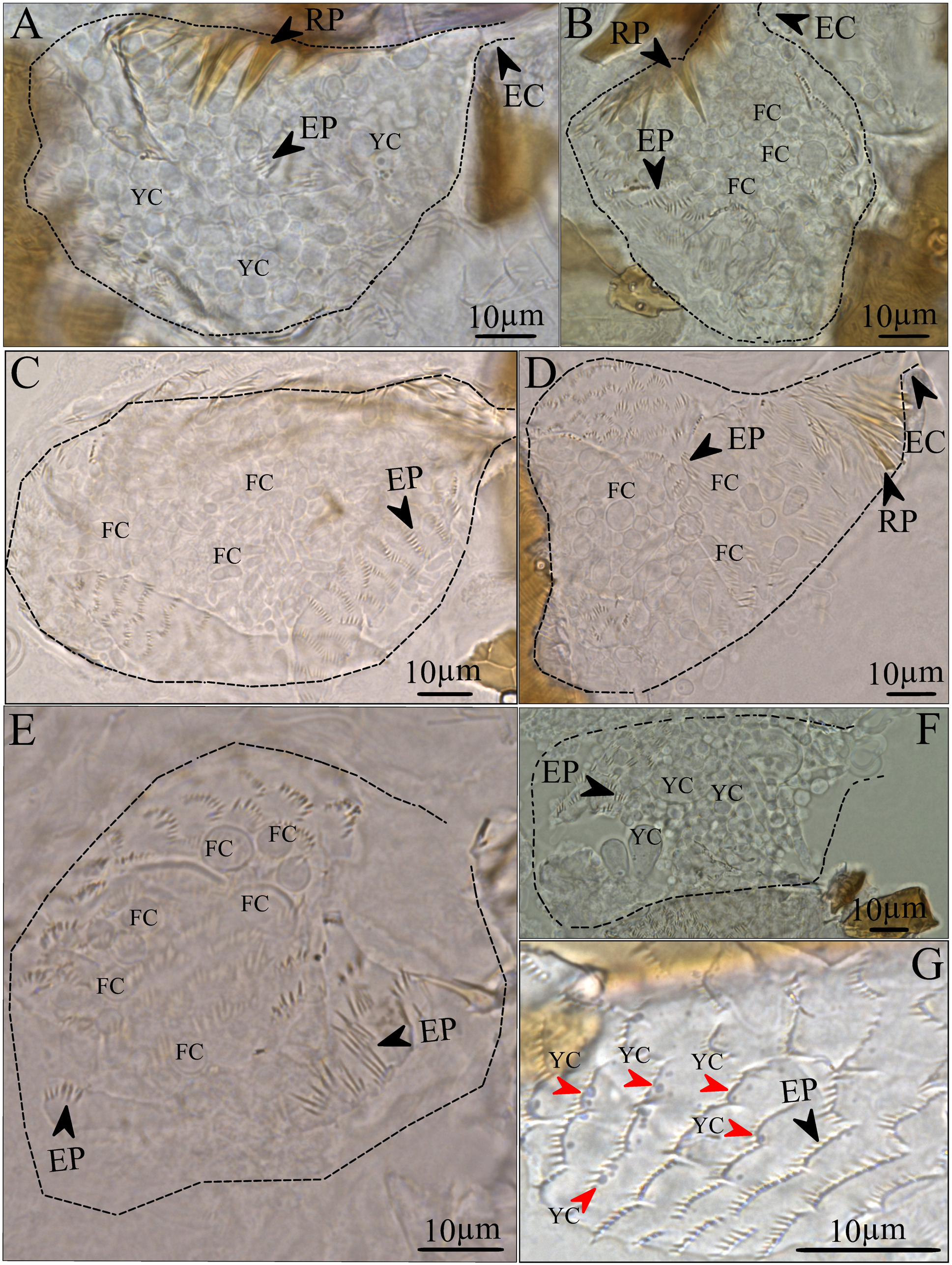
Light micrographs of *X. affinis* mycangial sections after colonization by indicated fungal and yeast symbionts. The mycangial boundaries are indicated by dashed lines. Structural features include Ross projections (RP), eyelash projections (EP), and entry channels (EC). Fungal cells (FC) and yeast cells (YC) are labeled within each panel. Mycangia colonized by: (A) *Ambrosiozyma monospora*, (B) *Neocosmospora affinis*, (C) *Graphium ambrosium*, (D) *Raffaelea arxii*, (E) *Raffaelea fusca*, (F) *Alloascoidea xylebori*, and (G) *Wickerhamomyces ambrosius*, with individual yeast cells indicated by red arrowheads. Scale bars = 10 μm

## Discussion

The symbiotic association of ambrosia beetles with their fungal partners evolved independently within almost a dozen different beetle-fungal lineages, dates from 40-60 million years, and includes devastating plant pathogens (Ploetz et al., 2013; Hughes et al., 2017; Vanderpool et al., 2018). This mutualism has coincided with the evolution of specialized beetle organs, termed mycangia (Mayers et al., 2022). Between different beetle lineages, these structures vary tremendously in form, location, and mechanism of symbiont transport and selection. Morphological aspects of the mycangia of several beetle species, including several within the *Xyleborus* genera, have been examined (Jiang et al., 2019; Spahr et al., 2020), and identification of presumed beetle symbionts has included both culturing/isolation and metagenomic approaches (Bateman et al., 2016; Campbell et al., 2016; Skelton et al., 2019).

However, the missing information in almost all such studies is a demonstration that the identified fungi can then re-colonize and be re-isolated from the mycangia of the hosts from which they were originally identified. Koch’s postulates, designed to establish causation between a specific microbe and a disease, involves meeting four basic criteria, namely, demonstrating; (i) association: the microbe must be found in organisms suffering from the disease, but is either absent or at low levels in healthy organisms, (ii) isolation: the microbe should be isolated from a diseased organism and grown in pure culture [note for many fastidious microbes this may not currently be possible]. (iii) infectivity: the microbe should cause the original disease when introduced into a healthy, susceptible host, and (iv) re-isolation: the microbe should be re-isolated from the experimentally infected host. With respect to symbionts (mutualists), this paradigm can be slightly altered to consider (i) identification of associated microbes, (ii) isolation, (iii) symbiosis establishment, i.e., in this case, the ability to colonize the mycangia, and (iv) re-isolation (from the mycangia). As noted, while the first two criteria have been widely met, confirmation of the latter has been lacking.

*X. affinis* is a widely distributed ambrosia beetle species, reported to be able to attack >245 different woody plant species, particularly problematic in sugar cane plantations, but affecting other economically important trees, e.g., mango and cacao amongst others (Biedermann, 2020; Rodriguez-Becerra et al., 2024). A variety of fungal species have been isolated from (Kostovcik et al., 2015). These included *Raffaelea* and *Fusarium* spp, *Cephalosporium pallidum*, and *Ambrosiella*, *Candida*, *Pischia*, and others, as well as unspecified filamentous fungi and yeasts. *H. lauricola* and *Raffaelea* spp. (*R. arxii*, *R. subalba*, and *R. subfusca*) were seen predominating in *Xyleborus* spp. with *R. arxii* the most abundant symbionts of *X. affinis*, as well as yeasts consistently found in their mycangia (Saucedo-Carabez et al., 2018). A meta-genomic barcoding study identified 40 fungal and 428 bacterial operational taxonomical units (OTUs), with *Raffaelea* and four ascomycetes yeasts considered to represent a “core” community (Ibarra-Juarez et al., 2020). Here, we isolated seven different fungal and yeast taxa from both *X. affinis* beetle gallery walls and from mycangia. Three corresponded to previously characterized *Xyleborus*-associated symbiotic fungi, i.e., *R. arxii*, *R. fusca*, and the yeast, *A. monospora*. The remaining four, we described as new species based on molecular phylogenetic, and morphological characteristics. These have been designated as: *N. affinis*, *G. ambrosium*, *A. xylebori*, and *W. ambrosius*; the former two are filamentous fungi, and the latter, yeasts. It should be noted that our analyses represent laboratory colonies, and hence a reflection of a more “core” community vertically transmitted, whereas the other reports indicated above have mainly focused on field capture, which is likely to be more diversified. Regarding mycangia enumeration, we (and others) routinely use an ethanol wash followed by water washes of the dissected head, before extraction (bead beating) and plating of symbionts to help minimize “background” fungal cells that may reside on the surface, within the mouthparts, and/or otherwise not in the mycangia (Harrington and Fraedrich, 2010; Ploetz et al., 2017; Saucedo et al., 2017; Saucedo-Carabez et al., 2018). However, we had noted significant variation in this method, and our structural studies suggested that the mycangia are likely to be open to the outside and that ethanol may enter and kill contents within the mycangia, thus biasing the results (Joseph et al., 2023; Joseph et al., 2025b; Joseph et al., 2025a). We, therefore, performed the extraction with and without the ethanol wash, and plated aliquots from the subsequent water washes to determine whether and/or the extent to which “background” fungal CFUs could be recovered under each protocol. These data revealed that CFU recovery in the last wash, irrespective of the ethanol wash, was typically <1 CFU/head, indicating that the ethanol wash was unnecessary. More importantly, total fungal recovery (CFUs) was higher after omitting the ethanol wash, and we were able to recover taxa previously missed, i.e., the new fungal species reported herein.

To fulfill the Koch-like postulates for symbioses as outlined above, we then demonstrated that all seven microbial taxa, as well as the previously reported *X. affinis* symbiont, *H. lauricola*, were competent in being able to colonize and be recovered from the mycangia, whereas the (non-mycangial) *Candida albicans* yeast could not colonize the mycangia, consistent with a previous study which showed that the fungal plant pathogen *Magnaporthe orzyae* also could not colonize

*X. affinis* mycangia (Joseph et al., 2023). These analyses further confirmed that all fungi showed decreased recovery after ethanol wash, with several symbionts, particularly the yeasts, being highly sensitive and would otherwise have been missed. More importantly, our results suggest a critical re-evaluation of ethanol washes in the enumeration of mycangial contents and the importance of plating (at the very least) the last water wash before extraction to confirm low “background” and the validity of the method. Intriguingly, we had expected that the revised (no-ethanol) wash might also reduce the wide variation previously observed in mycangial enumeration experiments (Joseph et al., 2023; Joseph et al., 2025b). Although overall recovery was significantly higher (with no ethanol wash), a significant degree of variation between individuals, i.e., some with apparently low or no colonization, continued to be seen across fungal taxa. We had previously speculated that this could be due to factors including mycangial dynamics, i.e., those with few fungi happened to be in the process of expelling their contents, to age or temporal competency, i.e., the mycangia were “closed” for some reason in those beetles, and/or to beetle feeding behaviors resulting in fungi not being exposed to the mycangia in the first place. Overall, these findings suggest that there are important aspects of mycangial functioning which require further investigation.

Our data also show that the dynamics and overall levels of fungal cells recovered from mycangia differed across the taxa tested, with some taxa colonizing the mycangia within 24 h and remaining stable over time (to 48 h), and others, e.g., *W. ambrosius*, found in the later time point. All fungal taxa grew apparently as yeast-like forms within the mycangia, with little to no hyphal growth seen. Qualitative imaging of the mycangia was congruent with the quantitative enumeration and, particularly for the yeasts, looked similar to *in vitro* grown cells. Notably, (i) *A. monospora*, *N. affinis*, *G. ambrosium, R. arxii*, and *A. xylebori* showed robust, i.e., in terms of apparent cell numbers, colonization, with cells mainly circular (or rod-shaped for *G. ambrosium*), with the exception of *A. xylebori* that showed two discrete cell types, (ii) *R. fusca* showed less “packed” mycangia, producing small circular yeast-like cells, and (iii) few *W. ambrosius* cells could be seen in mycangial sections. For all isolates, the fungal cells in the mycangia appear to be “free-floating”, i.e., no clear attachment of the fungus to the walls or any structure within the mycangia, and, as previously mentioned, little to no filamentous growth. Many fungi, including important human pathogens such as *Candida albicans* and *Coccidioides* species, but also plant and insect pathogens, undergo dimorphic transitions within hosts, which facilitates their evasion of host defense responses (Sanchez-Martinez and Perez-Martin, 2001; Boyce and Andrianopoulos, 2015; Gauthier, 2015). All three filamentous fungal partners grew as yeast-like cells within the mycangia, suggesting that this transition is critical for successful colonization. Consistent with this, a recent transcriptomics analysis of *H. lauricola* colonization of *X. affinis* mycangia indicated suppression of genes involved in hyphal and filamentous growth, that coincided with distinct changes in patterns of gene expression as compared to cells growing in vitro or during fungal association with its plant host (Joseph et al., 2026). One limitation of the current study is that our experiments only consider each fungal isolate on its own, and that consortium, succession, and/or mixed taxa effects likely occur, affecting overall mycangial symbiont composition. Future work examining such next levels of interactions and their implications both in terms of beetle gallery development and mycangial colonization, as well as the contribution of individual fungi to the fitness and productivity of beetle colonies are warranted. Fungal-animal mutualisms remain significantly understudied yet represent important and consequential environmental and evolutionary systems. Our data provide a framework for treating ambrosia beetle-fungal symbioses as unique models for addressing a wide range of questions in microbial ecology and the evolution of cooperative systems.

## ACKNOWLEDGEMENTS

This work was supported in part by an NSF IOS-2418026 award to N.O.K.

## DATA AVAILABILITY STATEMENT

All data generated or analyzed during this study are included in this published article and its supplementary information files.

---

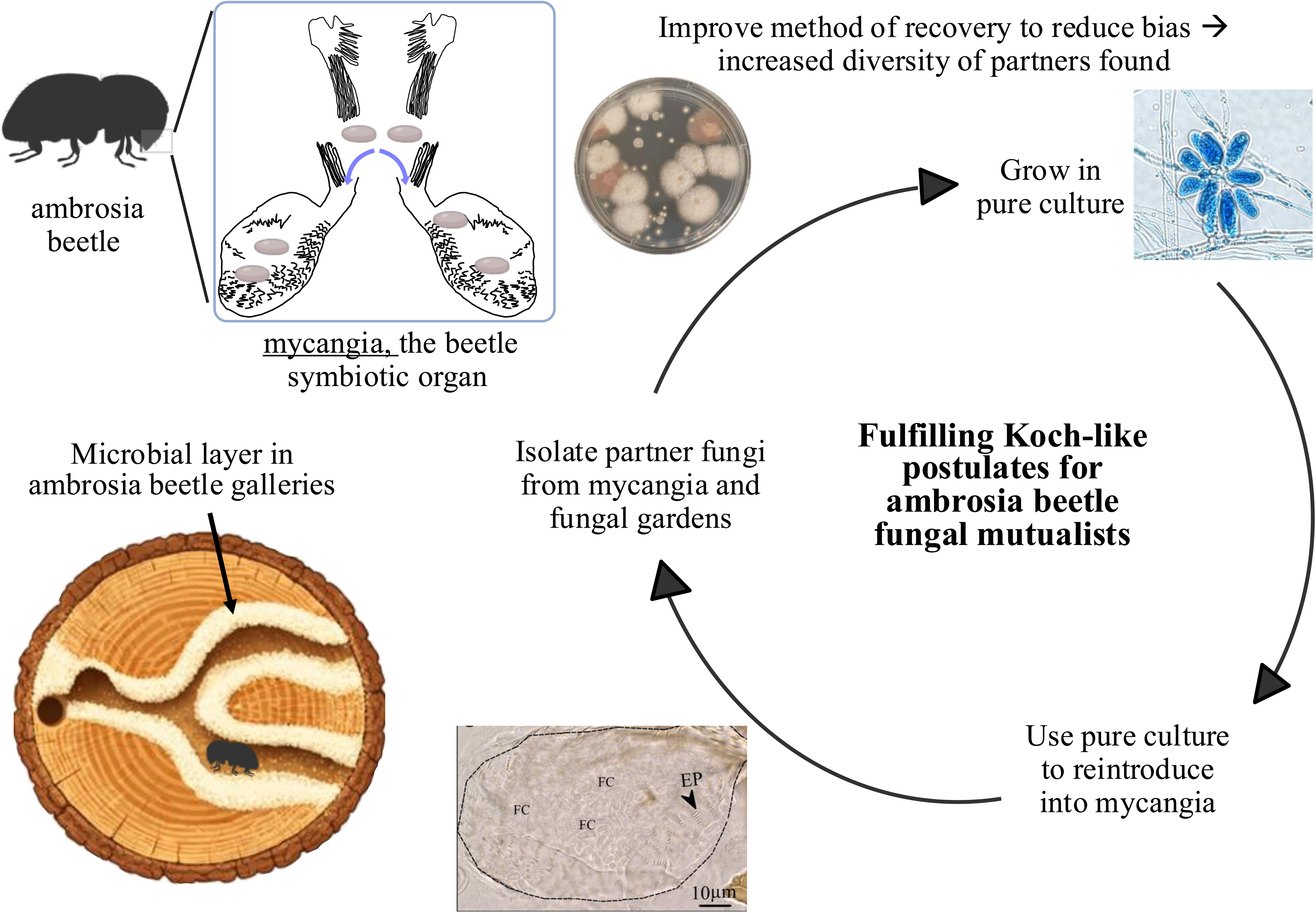

