## Supplementary Data for "Fulfilling Koch-like postulates for animal fungal mutualists: New fungal symbionts and gallery and mycangial colonization by *Xyleborus affinis* ambrosia fungi"

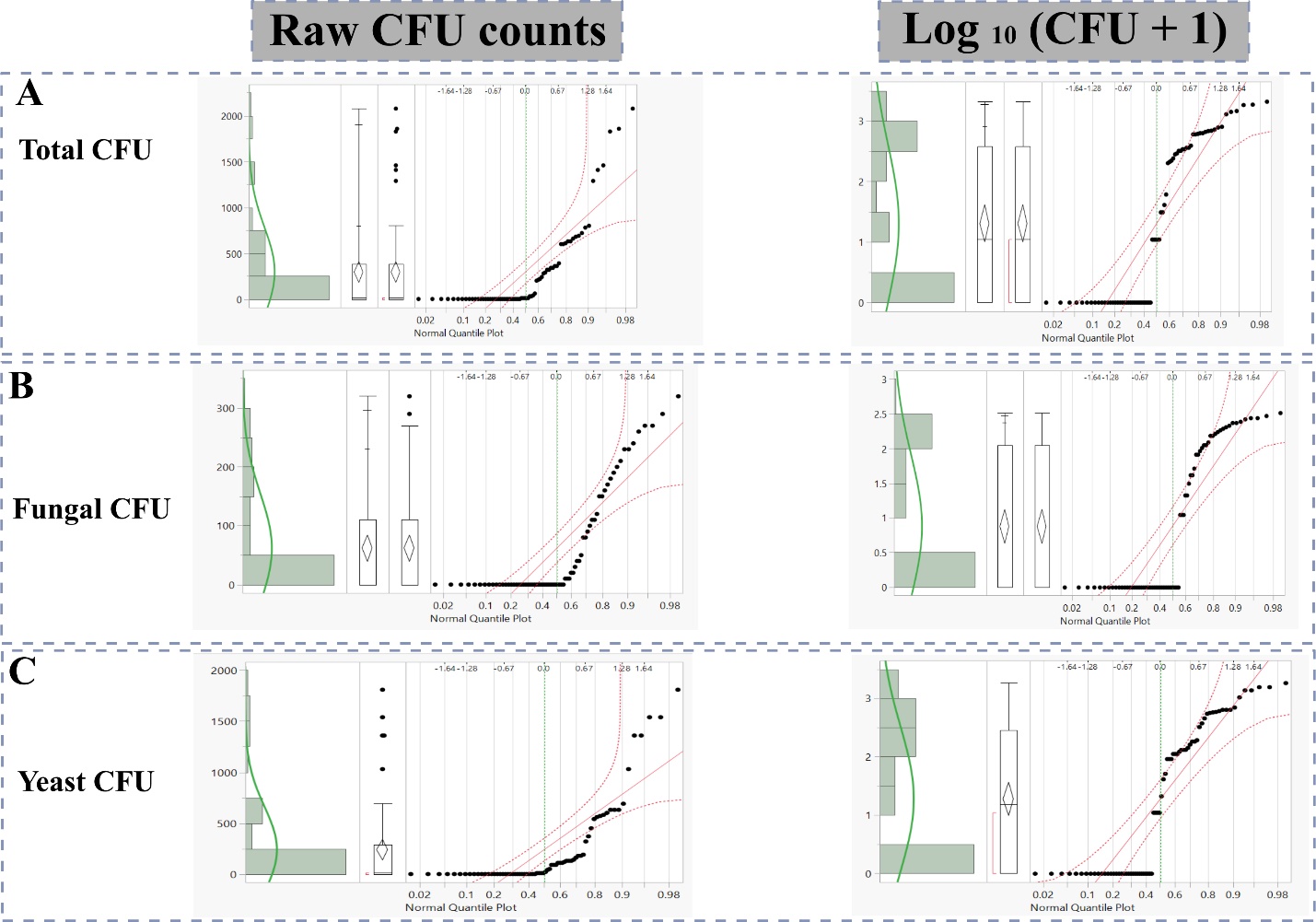

**Fig. S1**: Distribution and normality assessment of culturable microbial CFU counts obtained from *Xyleborus affinis* mycangia. Distributions of (A) total CFUs, (B) fungal CFUs, and (C) yeast CFUs are shown for raw CFU counts (left panels) and log_10_(CFU + 1)-transformed data (right panels). Each panel includes histograms with kernel density estimates, boxplots showing the median and interquartile range, and normal quantile-quantile (Q-Q) plots with 95% confidence envelopes. Raw CFU data exhibit strong right skewness and zero inflation, deviating from normality, whereas the log_10_(CFU + 1) transformation reduces skewness and improves symmetry but does not fully eliminate departures from normality.

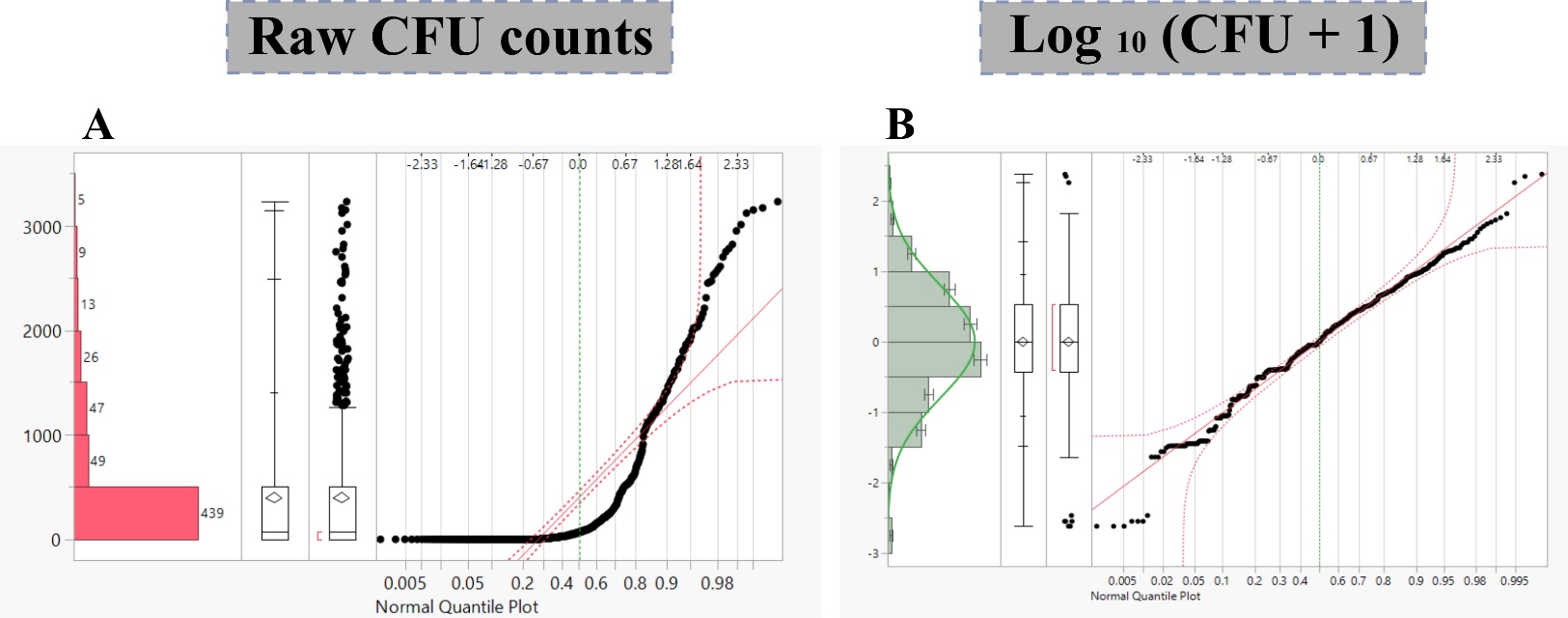

**Fig. S2**: Normality assessment of total CFU counts obtained from mycangial colonization bioassays. (A) Distribution of raw total CFU counts per mycangium, showing pronounced right skewness and zero inflation. (B) Distribution of log_10_ (CFU + 1)-transformed data, illustrating improved symmetry relative to the raw data. Each panel includes histograms with kernel density estimates, boxplots showing median and interquartile range, and normal quantile (Q-Q) plots with 95% confidence envelopes. Deviations from the theoretical normal distribution indicate departures from normality.

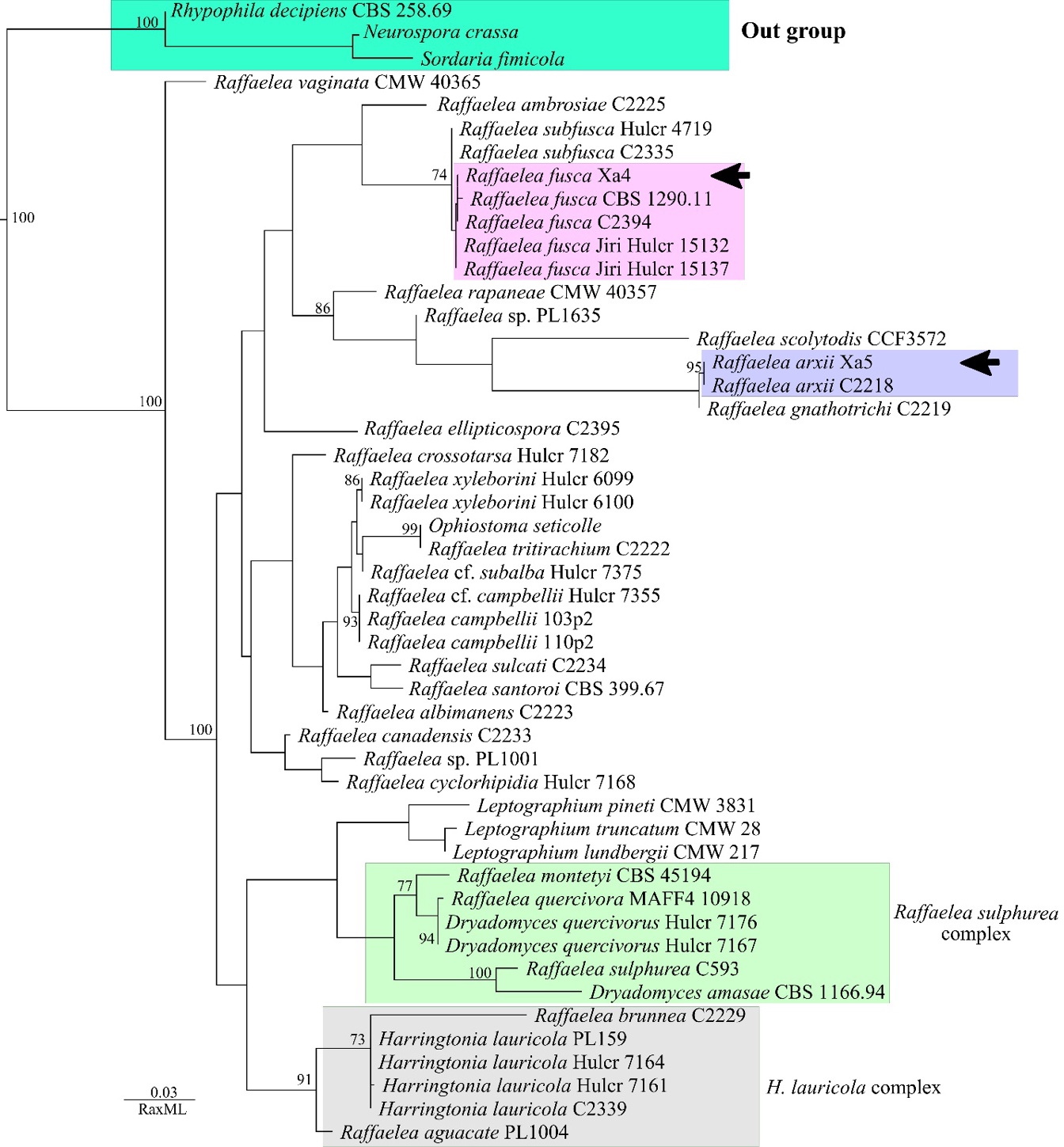

**Fig. S3**: Maximum likelihood phylogenetic tree of *Raffaelea* and related taxa inferred from LSU sequences using RAxML. Bootstrap support values (≥ 70%) are shown at the nodes. The tree is rooted with *Rhypophila decipiens*, *Neurospora crassa*, and *Sordaria fimicola* as outgroups (highlighted in turquoise). Major species complexes are indicated by shaded boxes, including the *Raffaelea sulphurea* complex (green) and the *Harringtonia lauricola* complex (gray). Isolates obtained in this study are highlighted in colored boxes, with *Raffaelea fusca* isolates shown in pink and *Raffaelea arxii* isolates shown in blue; arrows indicate representative strains discussed in the text. The scale bar represents substitutions per site.

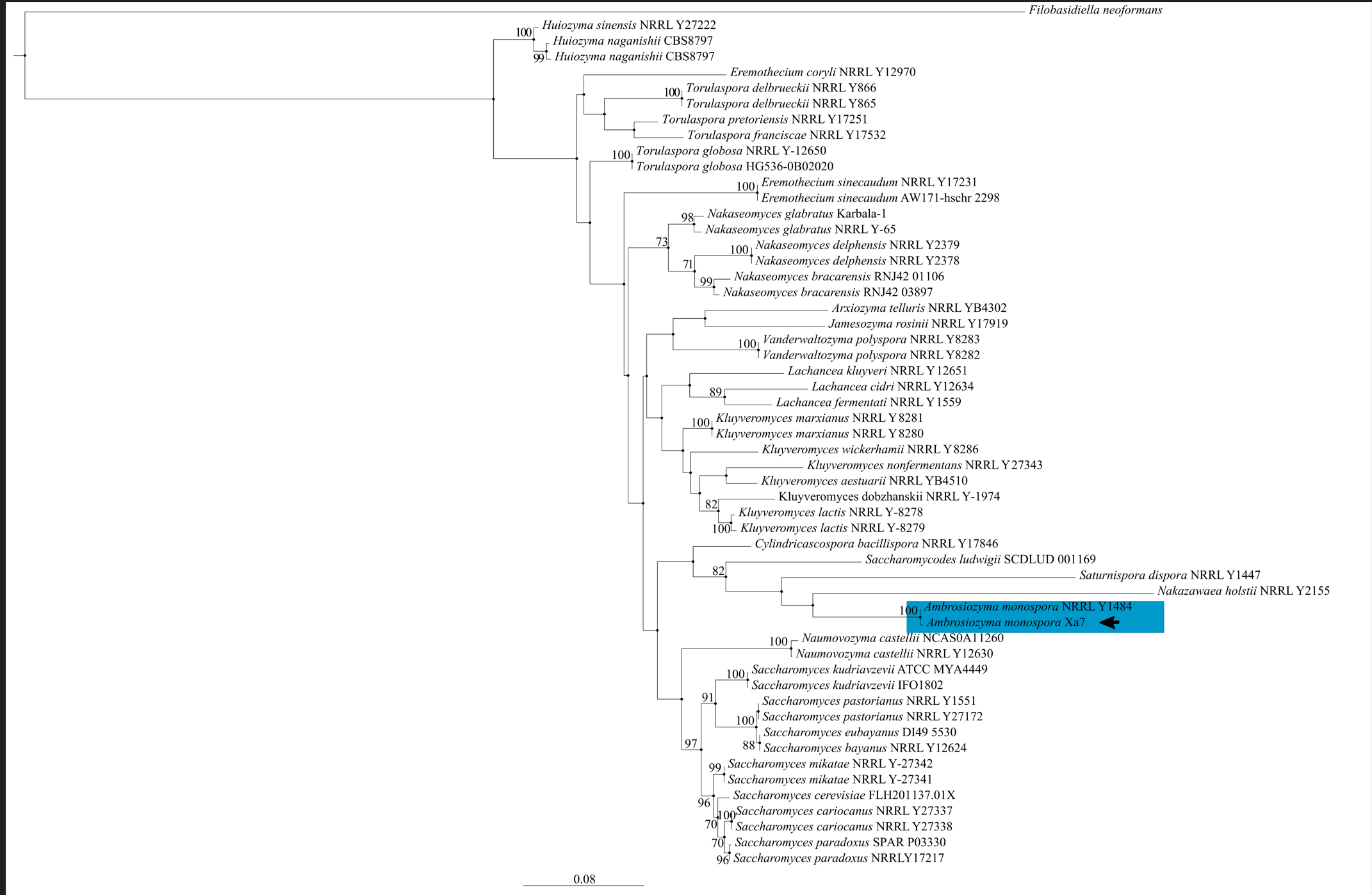

**Fig. S4**: Maximum likelihood phylogenetic tree based on *TEF* sequences showing the placement of *Ambrosiozyma monospora* relative to representative taxa within the Saccharomycetales. The tree was inferred using RAxML, and bootstrap support values ≥ 70% are shown at the nodes. The isolate obtained in this study (*A. monospora*) is highlighted in blue and indicated by an arrow. The tree is rooted with *Filobasidiella neoformans* as the outgroup. The scale bar represents substitutions per site.

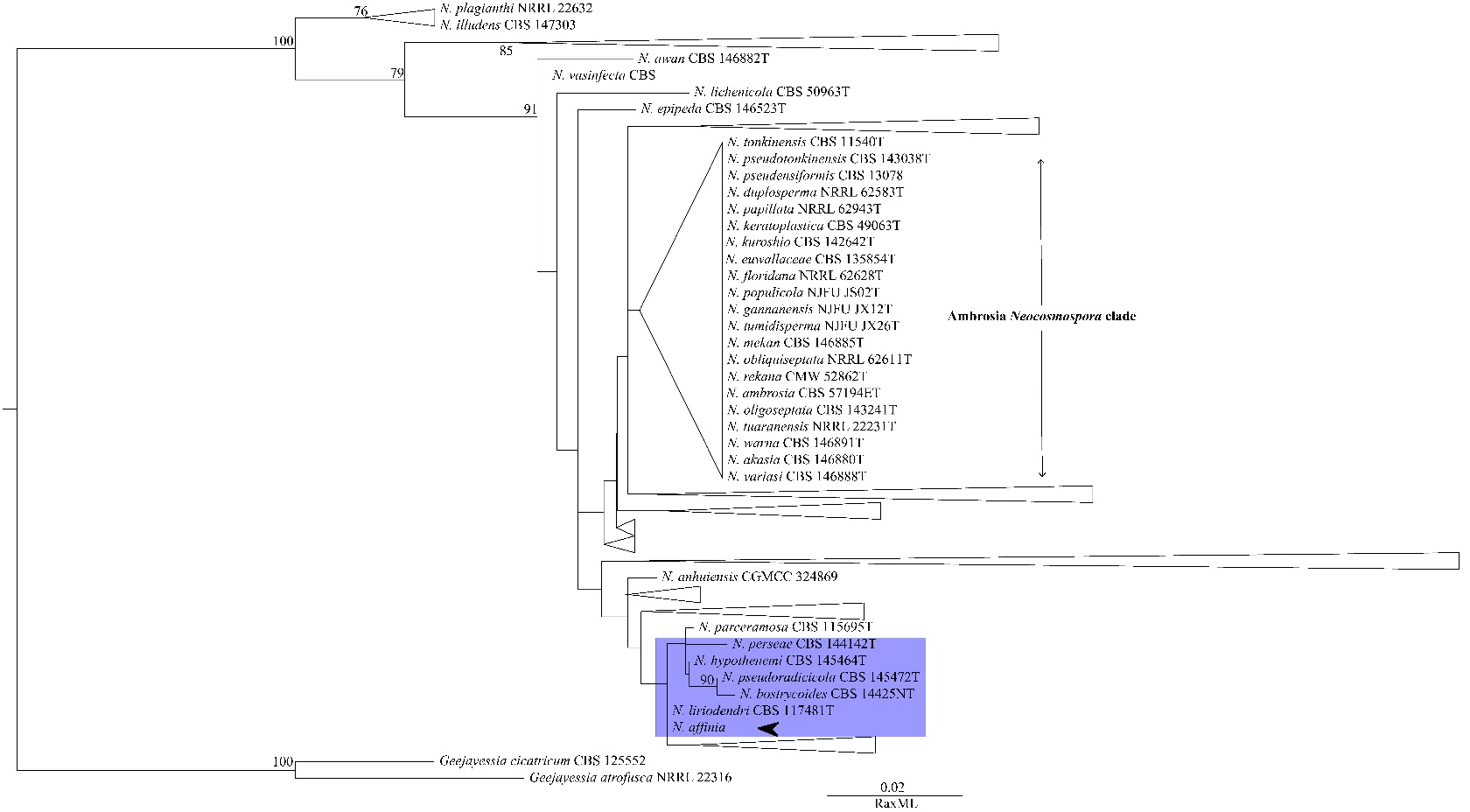

**Fig. S5**: Phylogenetic placement of *Neocosmospora affinis* inferred from *ITS* sequences. Maximum likelihood phylogenetic tree constructed using ITS rDNA sequences and inferred with RAxML. Bootstrap support values ≥ 70% are shown at the nodes. *Neocosmospora affinis* is highlighted and indicated by an arrow. *Geejayessia cicatricum* and *G. atrofusca* were used as outgroups. The scale bar represents the number of substitutions per site.

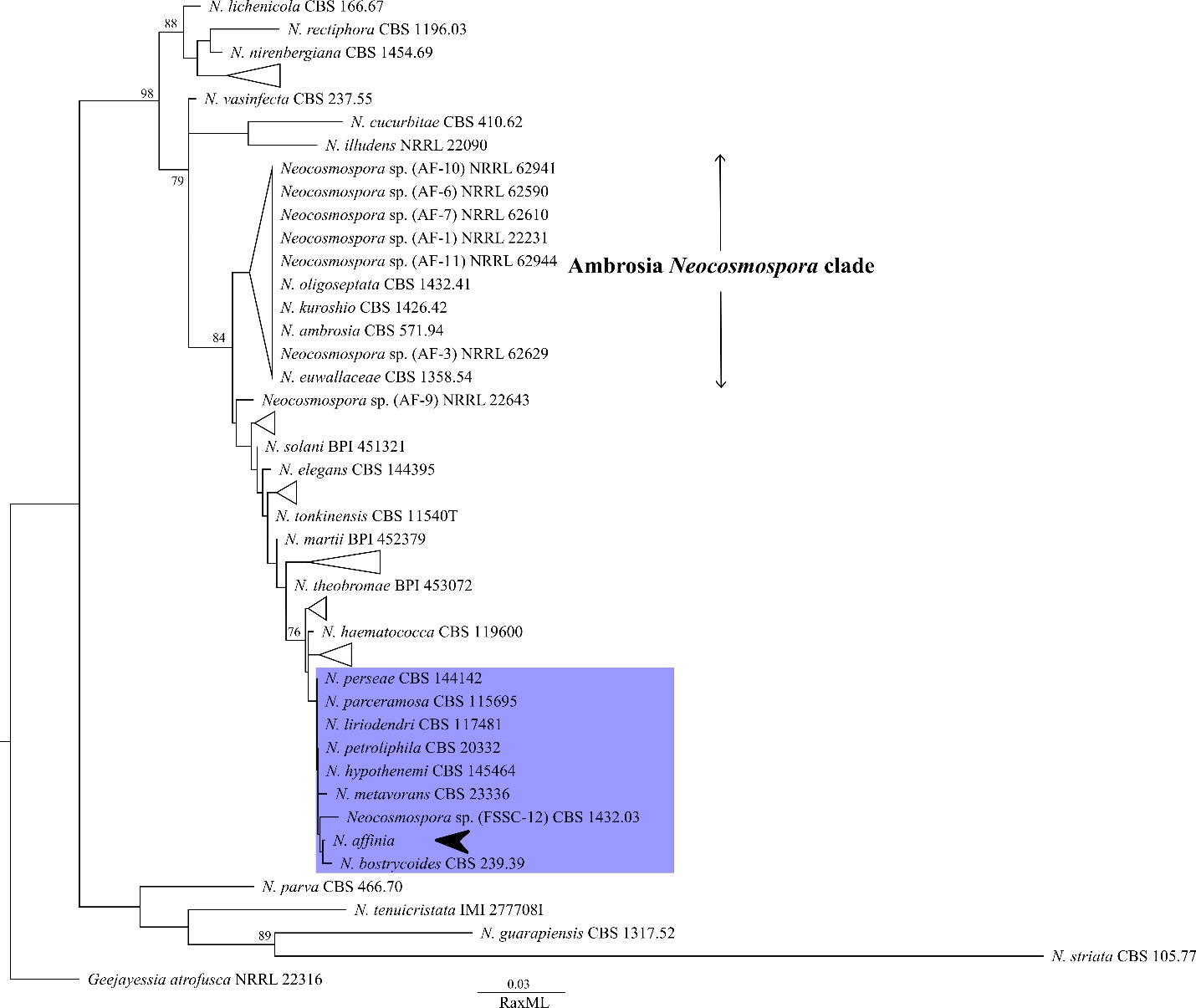

**Fig. S6**: Phylogenetic placement of *Neocosmospora affinis* inferred from LSU rDNA sequences. Maximum likelihood phylogenetic tree constructed using LSU rDNA sequences and inferred with RAxML. Bootstrap support values ≥ 70% are shown at the nodes. *Neocosmospora affinis* is highlighted and indicated by an arrow. *Geejayessia atrofusca* was used as the outgroup. The scale bar represents the number of substitutions per site.

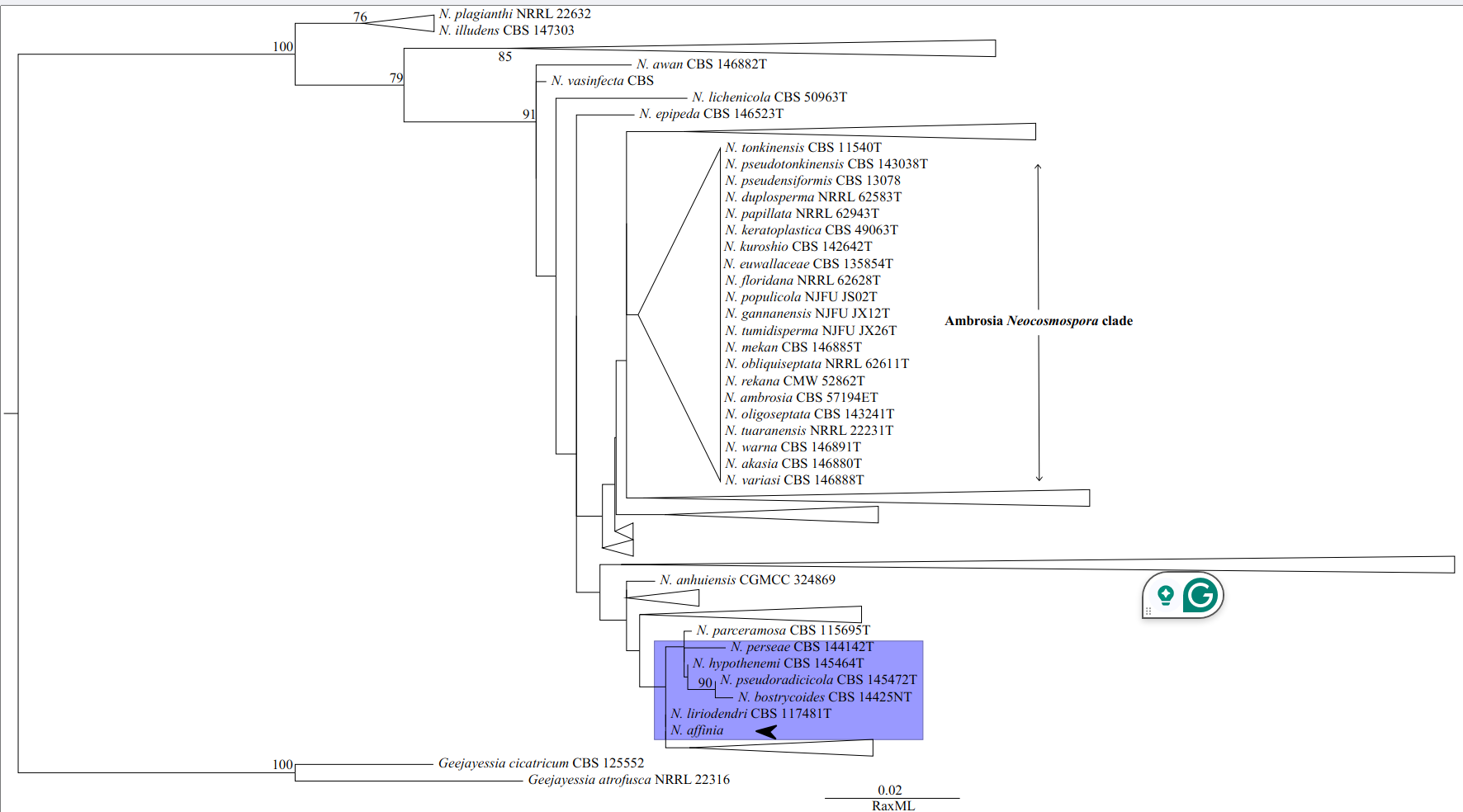

**Fig. S7**: Phylogenetic placement of *Neocosmospora affinis* inferred from *TEF1*-α sequences. Maximum likelihood phylogenetic tree constructed using partial *TEF1*-α sequences and inferred with RAxML. Bootstrap support values ≥ 70% are shown at the nodes. *Neocosmospora affinis* is highlighted and indicated by an arrow. *Geejayessia cicatricum* and *G. atrofusca* were used as outgroups. The scale bar represents the number of substitutions per site.

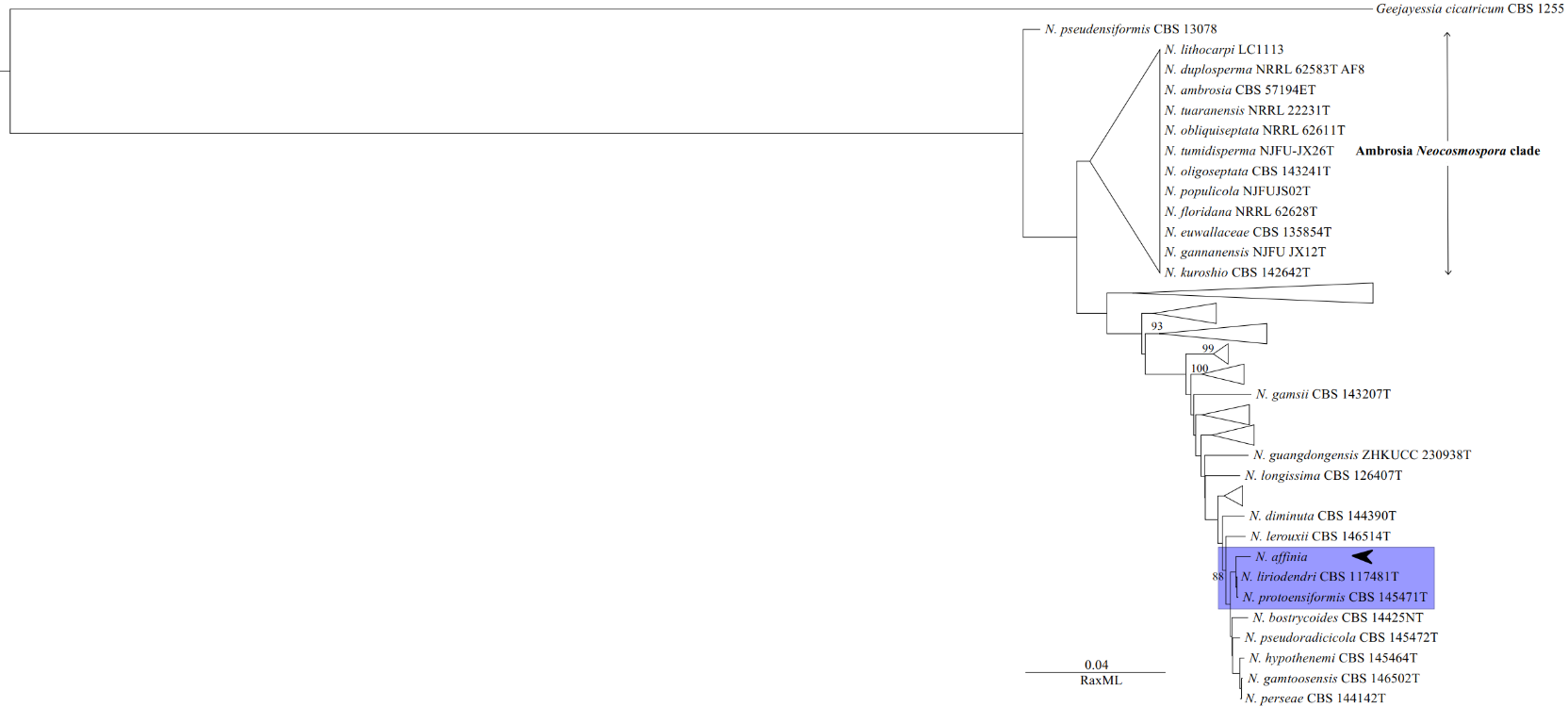

**Fig. S8**: Phylogenetic placement of *Neocosmospora affinis* inferred from *RPB1* sequences. Maximum likelihood phylogenetic tree constructed using partial *RPB1* sequences and inferred with RAxML. Bootstrap support values ≥ 70% are shown at the nodes. *Neocosmospora affinis* is highlighted and indicated by an arrow. *Geejayessia cicatricum* was used as the outgroup. The scale bar represents the number of substitutions per site.

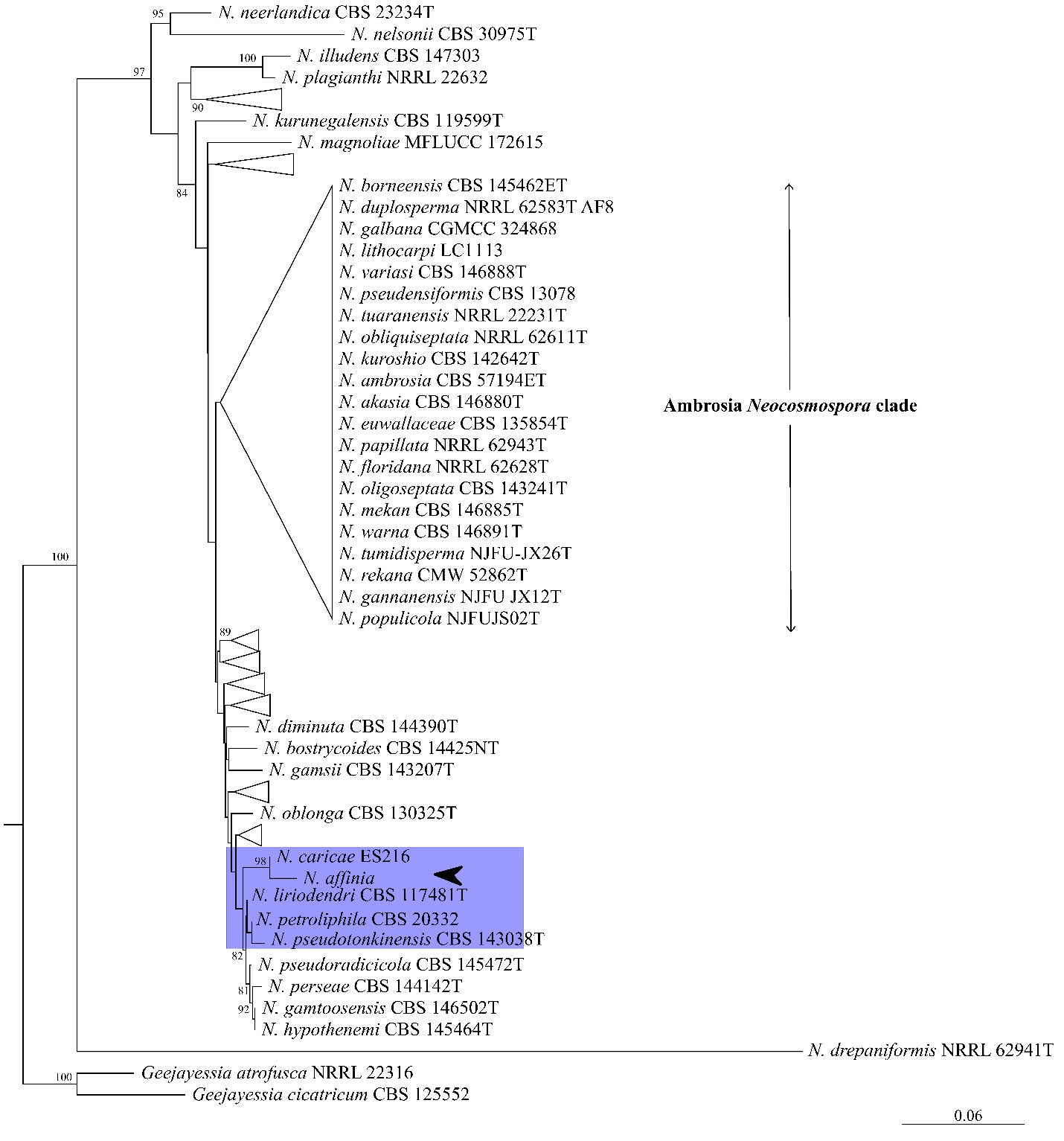

**Fig. S9**: Phylogenetic placement of *Neocosmospora affinis* inferred from *RPB2* sequences. A maximum-likelihood phylogenetic tree was constructed from partial *RPB2* gene sequences and inferred with RAxML. Bootstrap support values ≥ 70% are shown at the nodes. *Neocosmospora affinis* is highlighted and indicated by an arrow. The Ambrosia Neocosmospora clade is indicated. *Geejayessia atrofusca* and *G. cicatricum* were used as outgroups. The scale bar represents the number of substitutions per site.

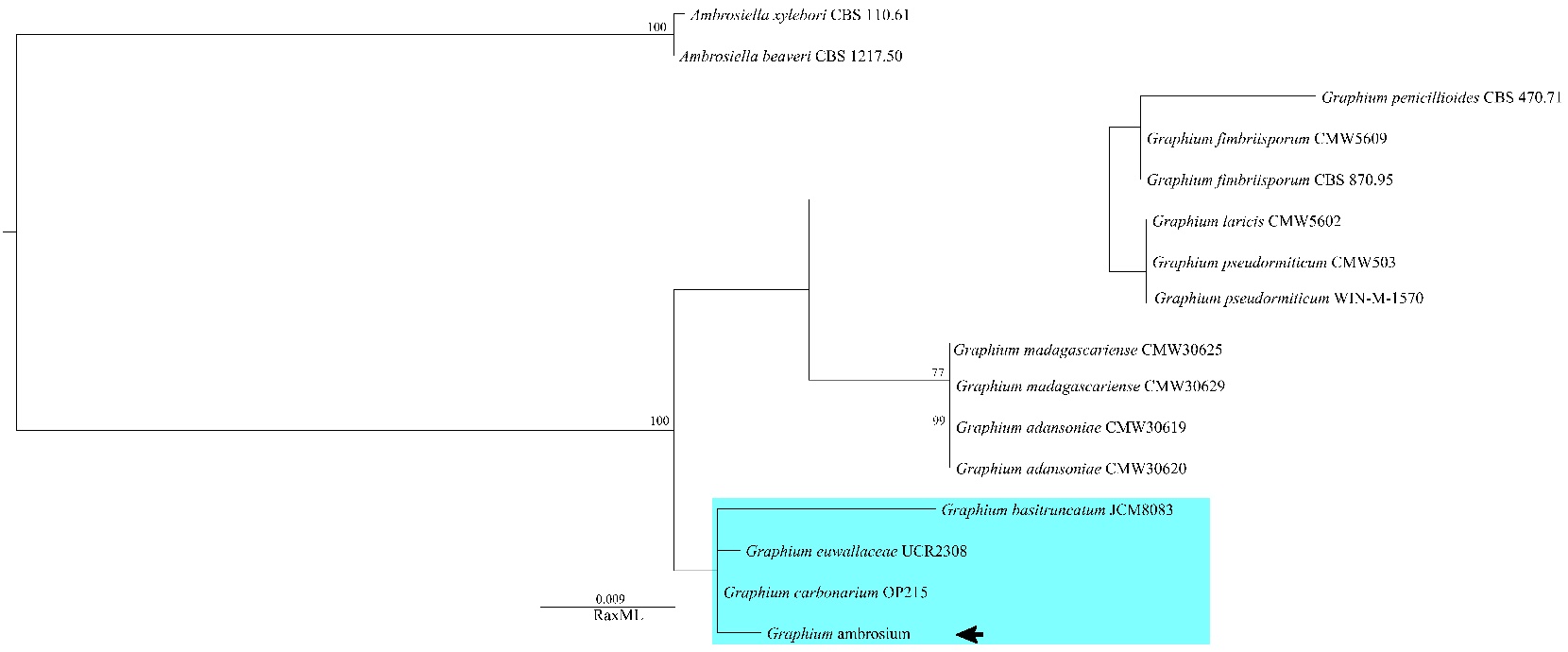

**Fig. S10**: Phylogenetic placement of *Graphium ambrosium* inferred from SSU rDNA sequences. Maximum likelihood phylogenetic tree constructed using SSU rDNA sequences and inferred with RAxML. Bootstrap support values ≥ 70% are shown at the nodes. *Graphium ambrosium* is highlighted and indicated by an arrow. *Ambrosiella xylebori* and *Ambrosiella beaveri* were used as outgroups. The scale bar represents the number of substitutions per site.

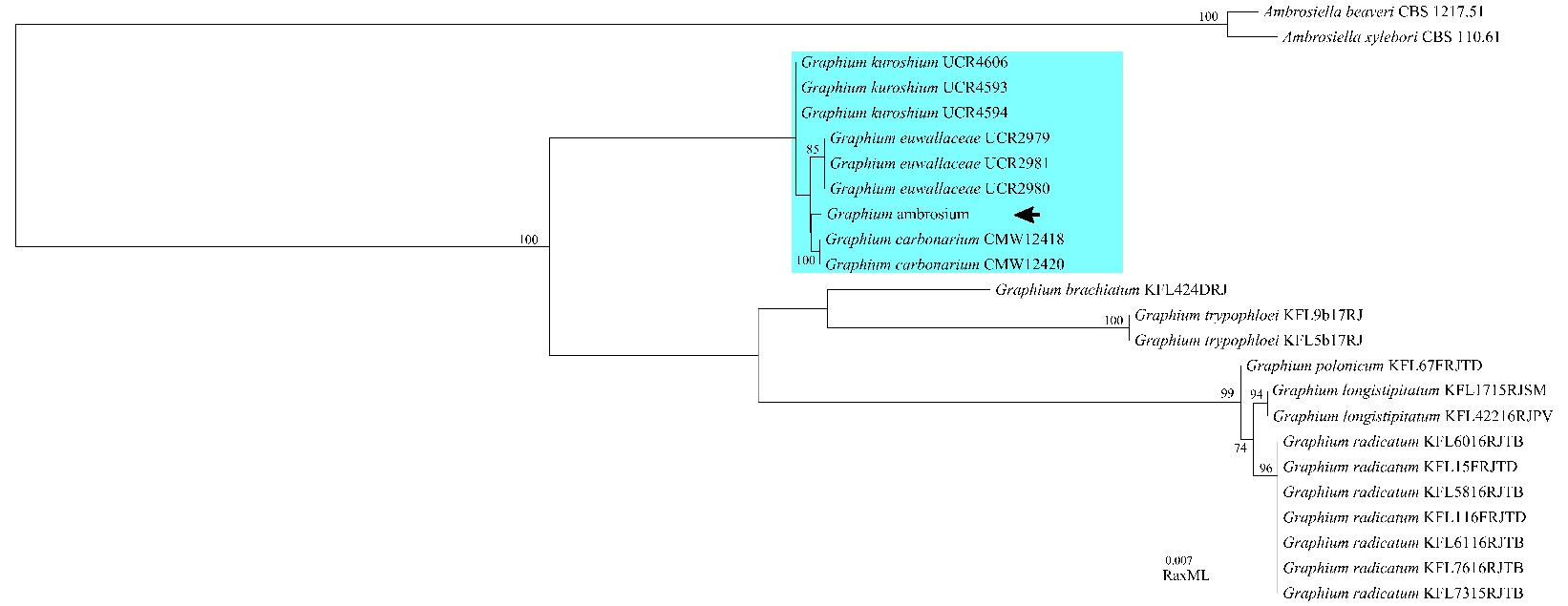

**Fig. S11**: Phylogenetic placement of *Graphium ambrosium* inferred from *β-tubulin* (*BTUB*) sequences. Maximum likelihood phylogenetic tree constructed using partial *β-tubulin* (*BTUB*) gene sequences and inferred with RAxML. Bootstrap support values ≥ 70% are shown at the nodes. *Graphium ambrosium* is highlighted and indicated by an arrow. *Ambrosiella xylebori* and *Ambrosiella beaveri* were used as outgroups. The scale bar represents the number of substitutions per site.

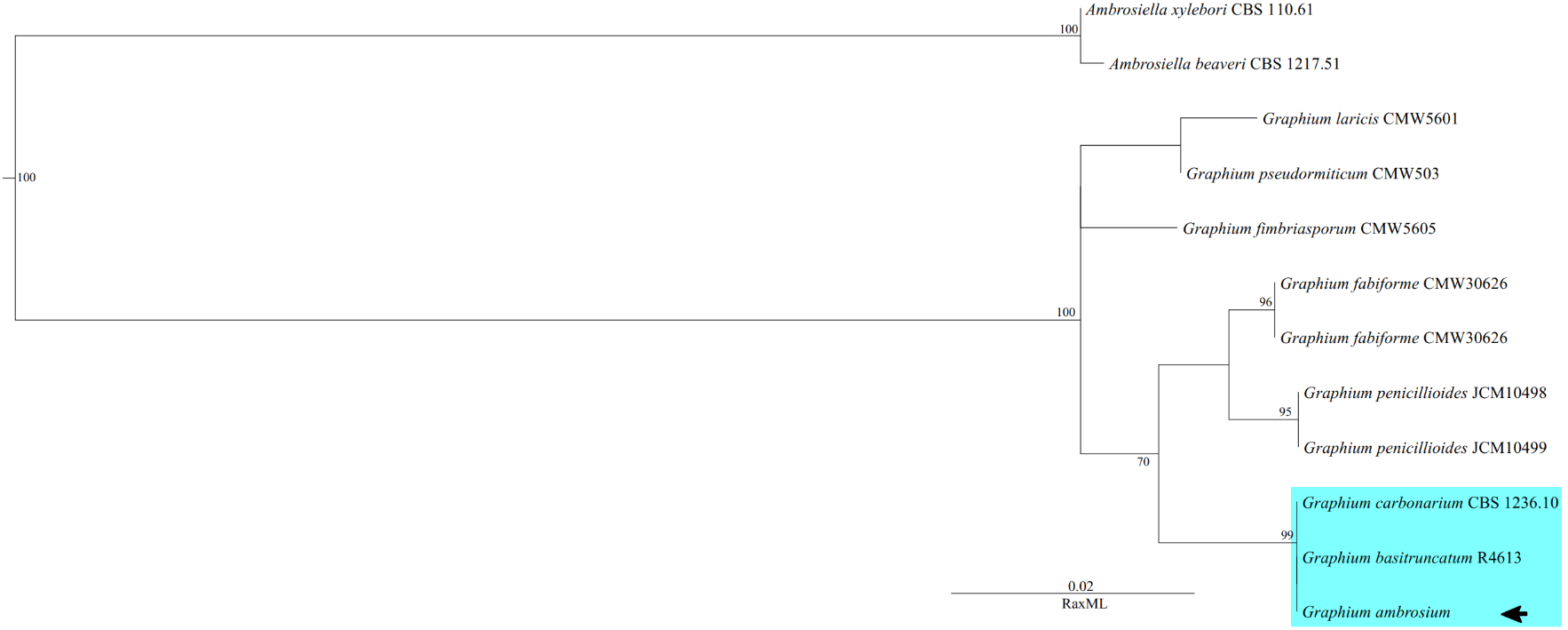

**Fig. S12**: Phylogenetic placement of *Graphium ambrosium* inferred from *LSU* rDNA sequences. Maximum likelihood phylogenetic tree constructed using *LSU* rDNA sequences and inferred with RAxML. Bootstrap support values ≥ 70% are shown at the nodes. *Graphium ambrosium* is highlighted and indicated by an arrow. *Ambrosiella xylebori* and *Ambrosiella beaveri* were used as outgroups. The scale bar represents the number of substitutions per site.

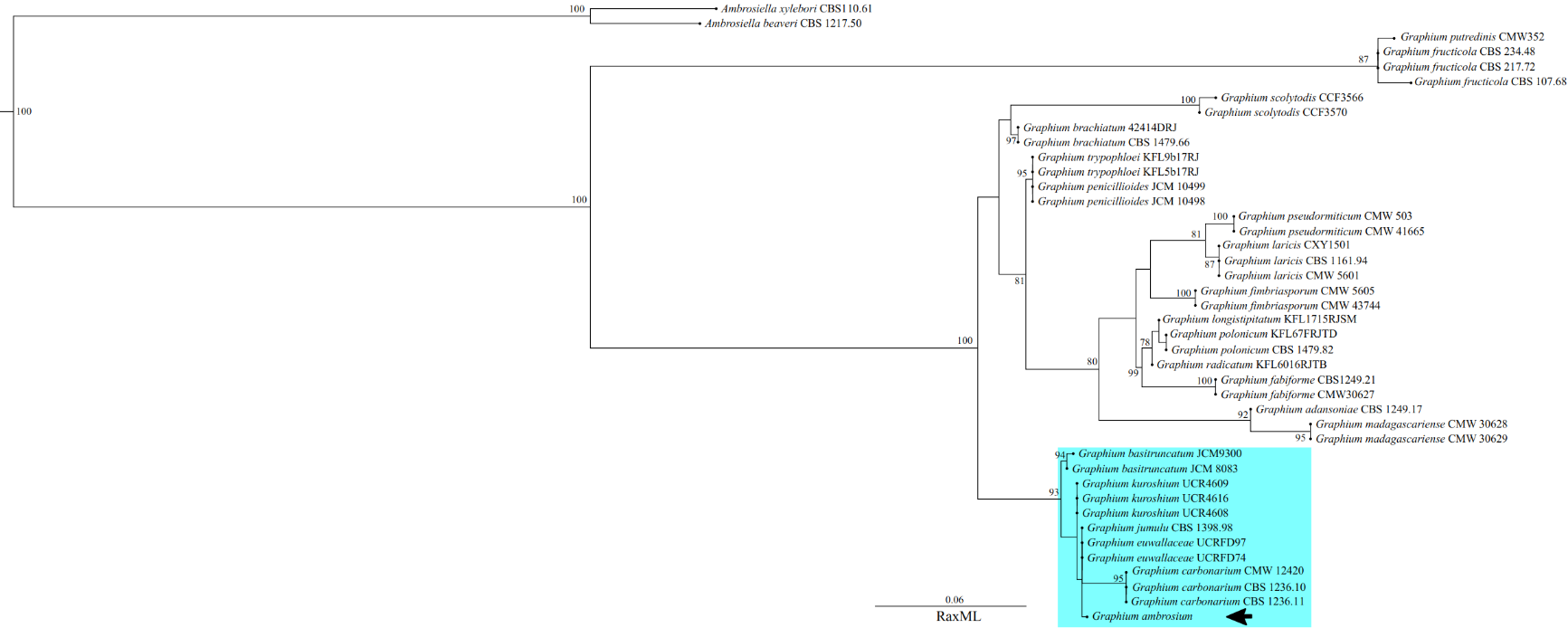

**Fig. S13**: Phylogenetic placement of *Graphium ambrosium* inferred from *ITS* rDNA sequences. Maximum likelihood phylogenetic tree constructed using *ITS* rDNA sequences and inferred with RAxML. Bootstrap support values ≥ 70% are shown at the nodes. *Graphium ambrosium* is highlighted and indicated by an arrow. *Ambrosiella xylebori* and *Ambrosiella beaveri* were used as outgroups. The scale bar represents the number of substitutions per site.

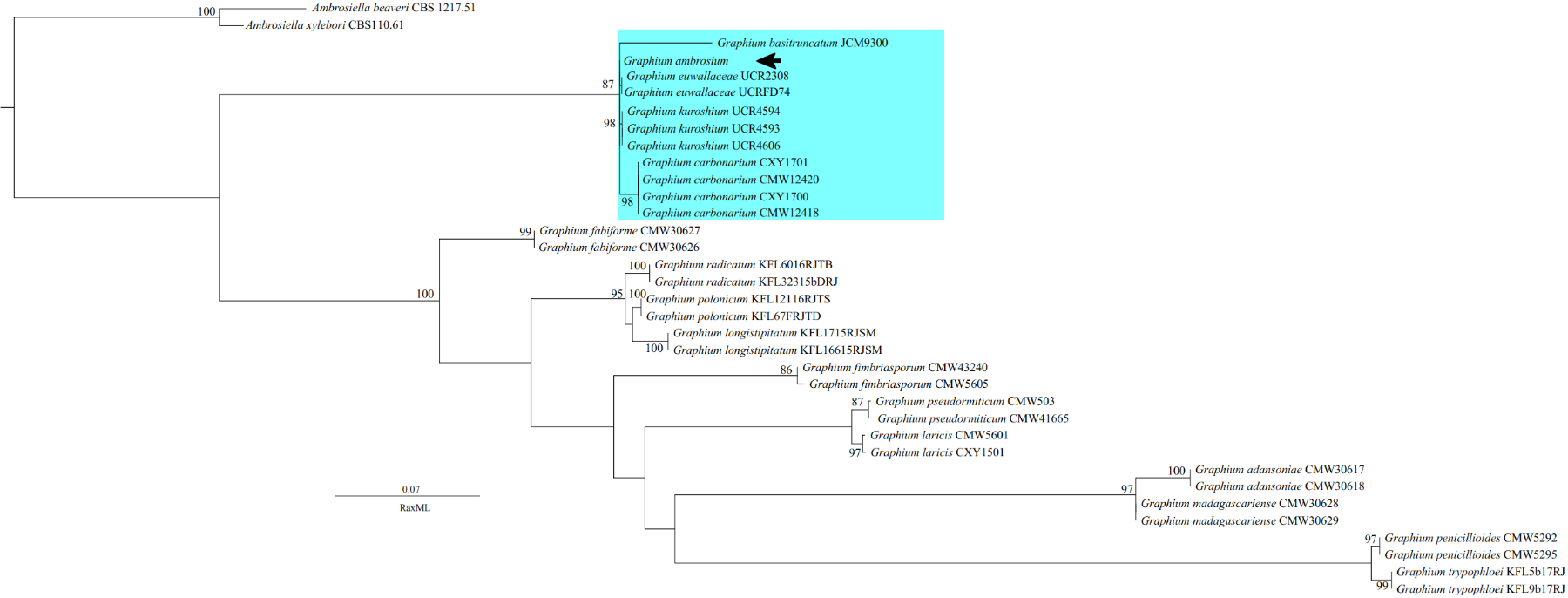

**Fig. S14**: Phylogenetic placement of *Graphium ambrosium* inferred from *TEF1*-α sequences. A maximum-likelihood phylogenetic tree was constructed from partial translation elongation factor 1-alpha (*TEF1*-α) gene sequences and inferred with RAxML. Bootstrap support values ≥70% are shown at the nodes. *Graphium ambrosium* is highlighted and indicated by an arrow. *Ambrosiella xylebori* and *Ambrosiella beaveri* were used as outgroups. The scale bar represents the number of substitutions per site.

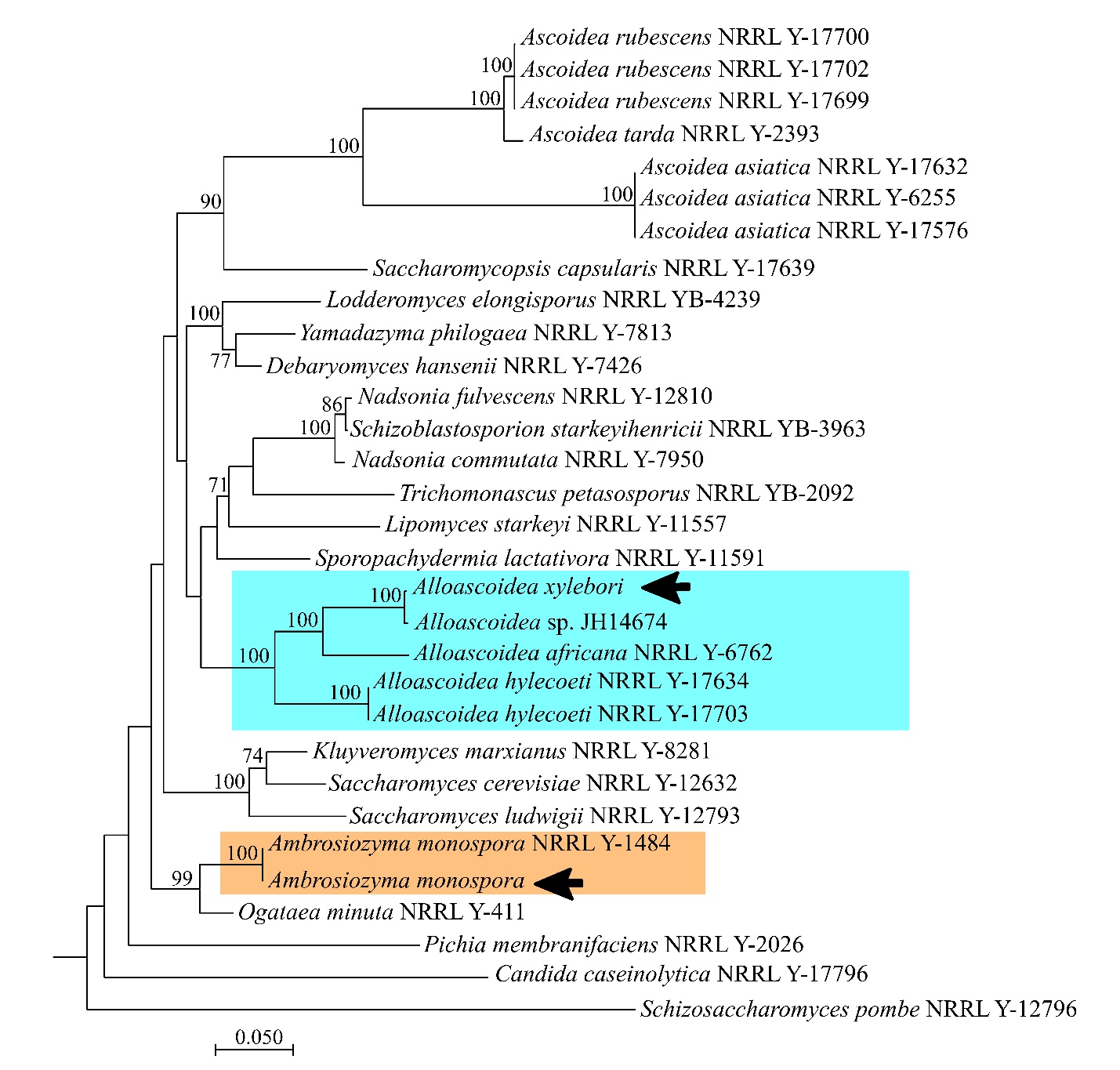

**Fig. S15**: Phylogenetic placement of *Alloascoidea xylebori* inferred from *LSU* rDNA sequences. Maximum likelihood phylogenetic tree constructed using *LSU* rDNA sequences and inferred with RAxML. Bootstrap support values ≥7 0% are shown at the nodes. *Alloascoidea xylebori* is highlighted and indicated by an arrow. The *Alloascoidea* clade is well supported, and *A. xylebori* clusters with *Alloascoidea* sp. JH14674, an isolate previously recovered from Florida. An isolate of *Ambrosiozyma monospora* generated in this study is included and highlighted. *Schizosaccharomyces pombe* was used as the outgroup. The scale bar represents the number of substitutions per site.

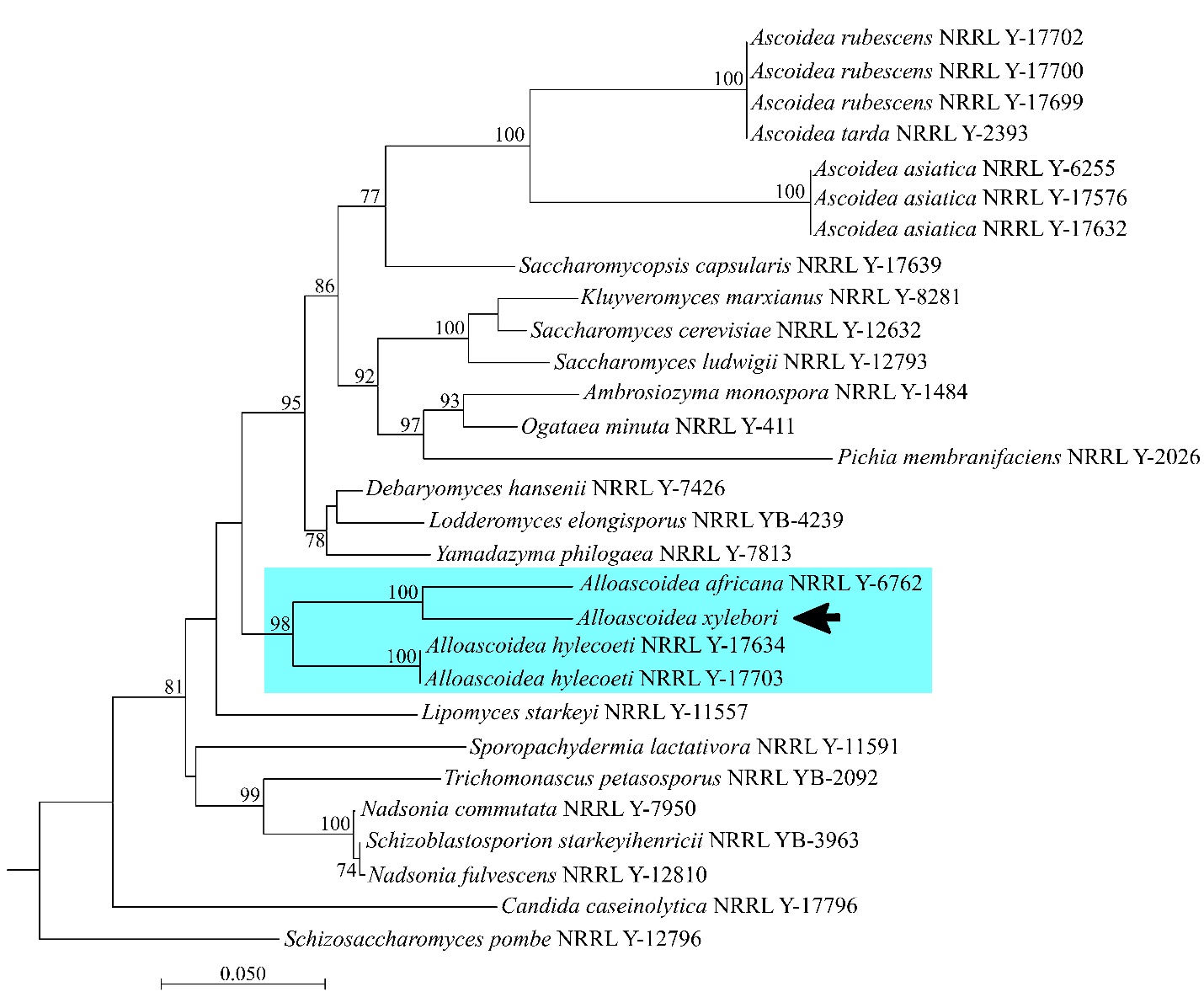

**Fig. S16**: Phylogenetic placement of *Alloascoidea xylebori* inferred from *SSU* rDNA sequences. Maximum likelihood phylogenetic tree constructed using *SSU* rDNA sequences and inferred with RAxML. Bootstrap support values ≥ 70% are shown at the nodes. *Alloascoidea xylebori* is highlighted and indicated by an arrow. *Schizosaccharomyces pombe* was used as the outgroup. The scale bar represents the number of substitutions per site.

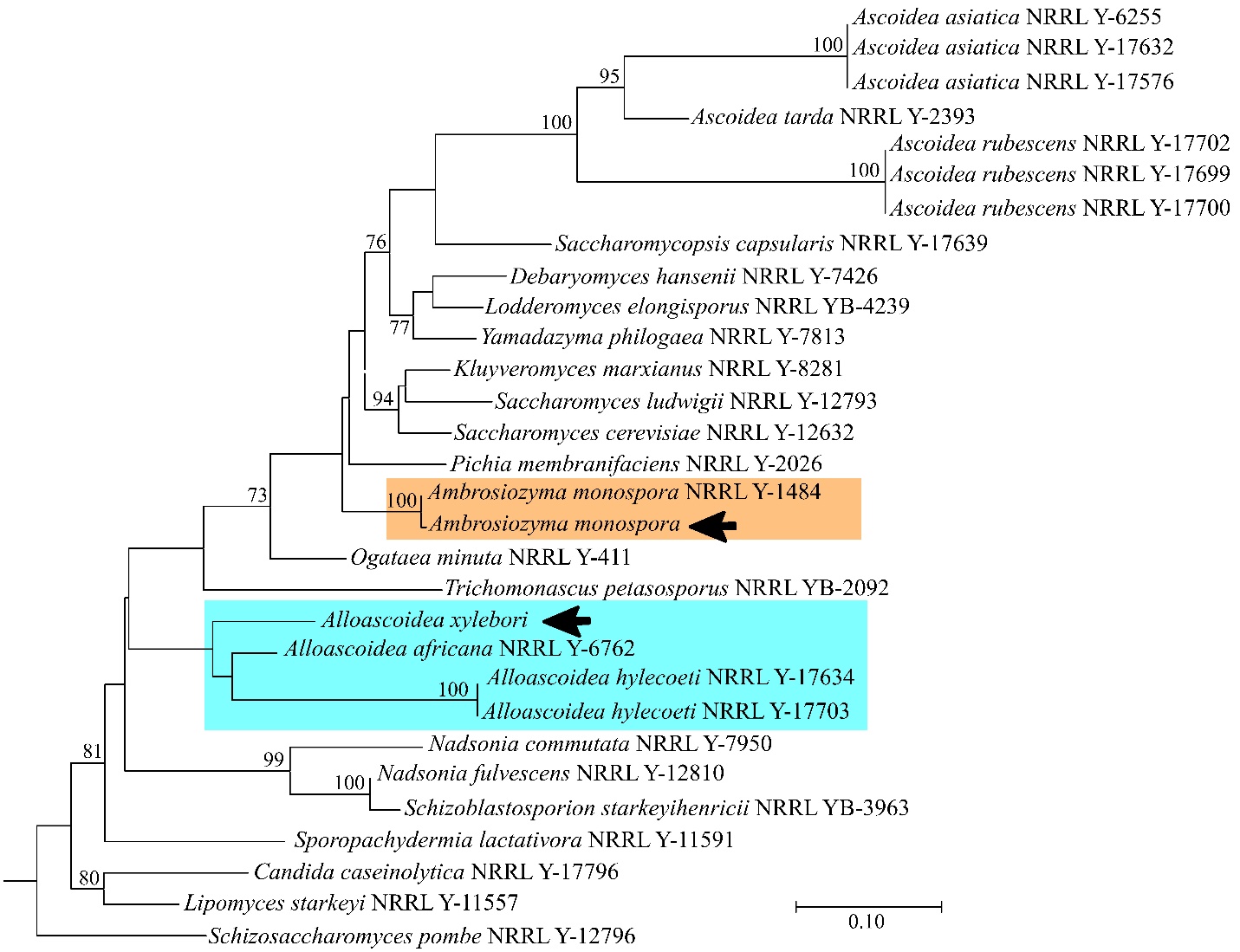

**Fig. S17**: Phylogenetic placement of *Alloascoidea xylebori* inferred from *TEF1*-α sequences. A maximum-likelihood phylogenetic tree was constructed from partial translation elongation factor 1-alpha (*TEF1*-α) gene sequences and inferred with RAxML. Bootstrap support values ≥ 70% are shown at the nodes. *Alloascoidea xylebori* is highlighted and indicated by an arrow, clustering within the *Alloascoidea* lineage. An isolate of *Ambrosiozyma monospora* generated in this study is included and highlighted for comparison. *Schizosaccharomyces pombe* was used as the outgroup. The scale bar represents the number of substitutions per site.

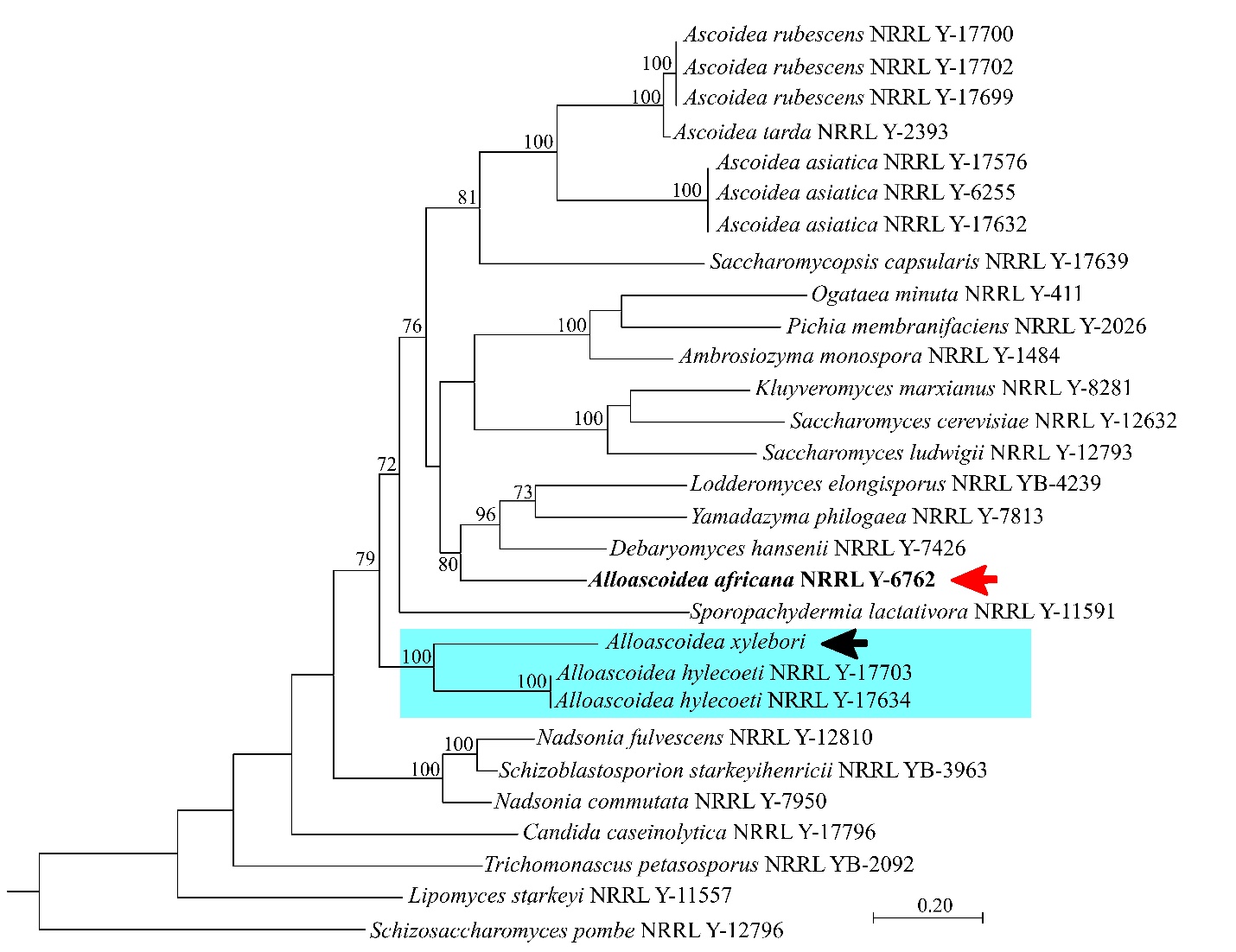

**Fig. S18**: Phylogenetic placement of *Alloascoidea xylebori* inferred from *RPB1* sequences. Maximum likelihood phylogenetic tree constructed using partial *RPB1* gene sequences and inferred with RAxML. Bootstrap support values ≥ 70% are shown at the nodes. *Alloascoidea xylebori* is highlighted and indicated by a black arrow. *Schizosaccharomyces pombe* was used as the outgroup. *Alloascoidea africana* (red arrow) is placed outside the main *Alloascoidea* cluster in this *RPB1* phylogeny; this placement is likely attributable to poor sequence quality, as the *RPB1* sequence contains a high proportion of ambiguous nucleotides (32 Ns), which may have affected its phylogenetic resolution. The scale bar represents the number of substitutions per site.

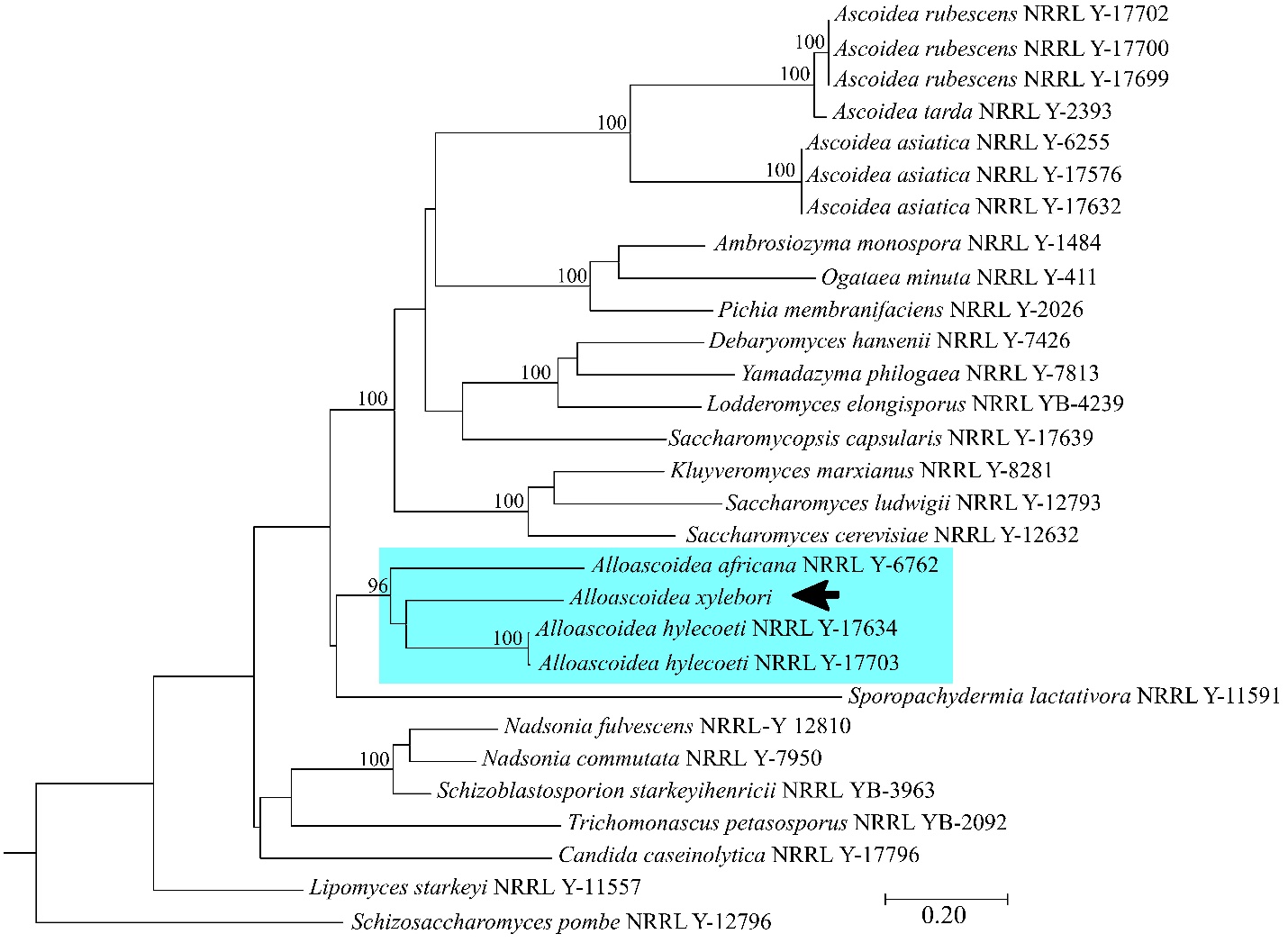

**Fig. S19**: Phylogenetic placement of *Alloascoidea xylebori* inferred from *RPB2* sequences. A maximum-likelihood phylogenetic tree was constructed from partial *RPB2* gene sequences and inferred with RAxML. Bootstrap support values ≥70% are shown at the nodes. *Alloascoidea xylebori* is highlighted and indicated by a black arrow*. Schizosaccharomyces pombe* was used as the outgroup. The scale bar represents the number of substitutions per site.

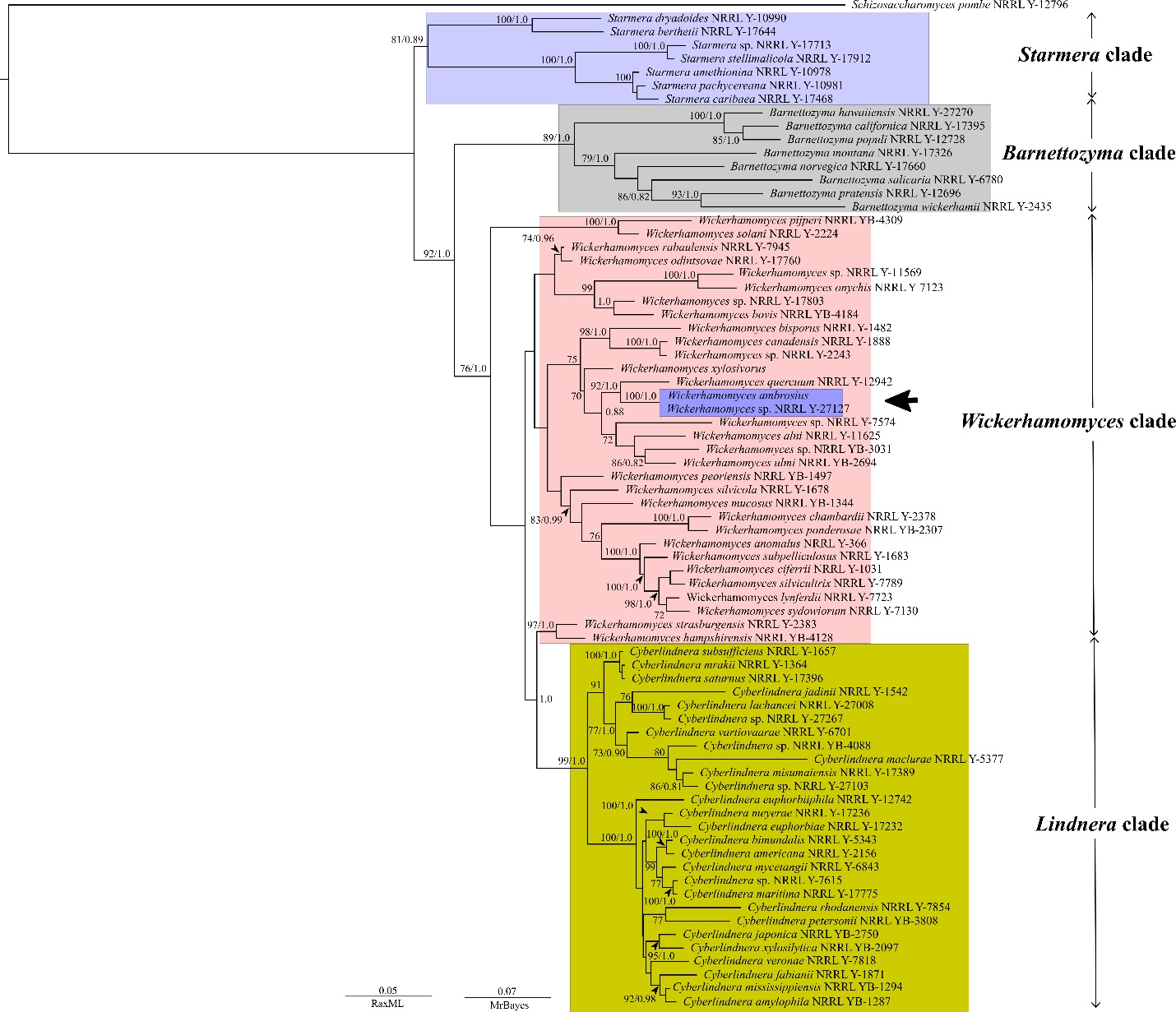

**Fig. S20**: Phylogenetic placement of *Wickerhamomyces ambrosius*. A concatenated maximum likelihood and Bayesian phylogenetic tree inferred from *ITS*, *LSU*, *SSU*, and *TEF* gene sequences. Numbers at nodes indicate bootstrap support values (RaxML) and posterior probabilities (MrBayes), respectively. The *Wickerhamomyces* clade is indicated, with *W. ambrosius* highlighted. Major related clades (e.g., *Barnettozyma*, *Lindnera*, and *Starmera*) are labeled for reference. *Schizosaccharomyces pombe* was used as the outgroup. The scale bar represents substitutions per site.

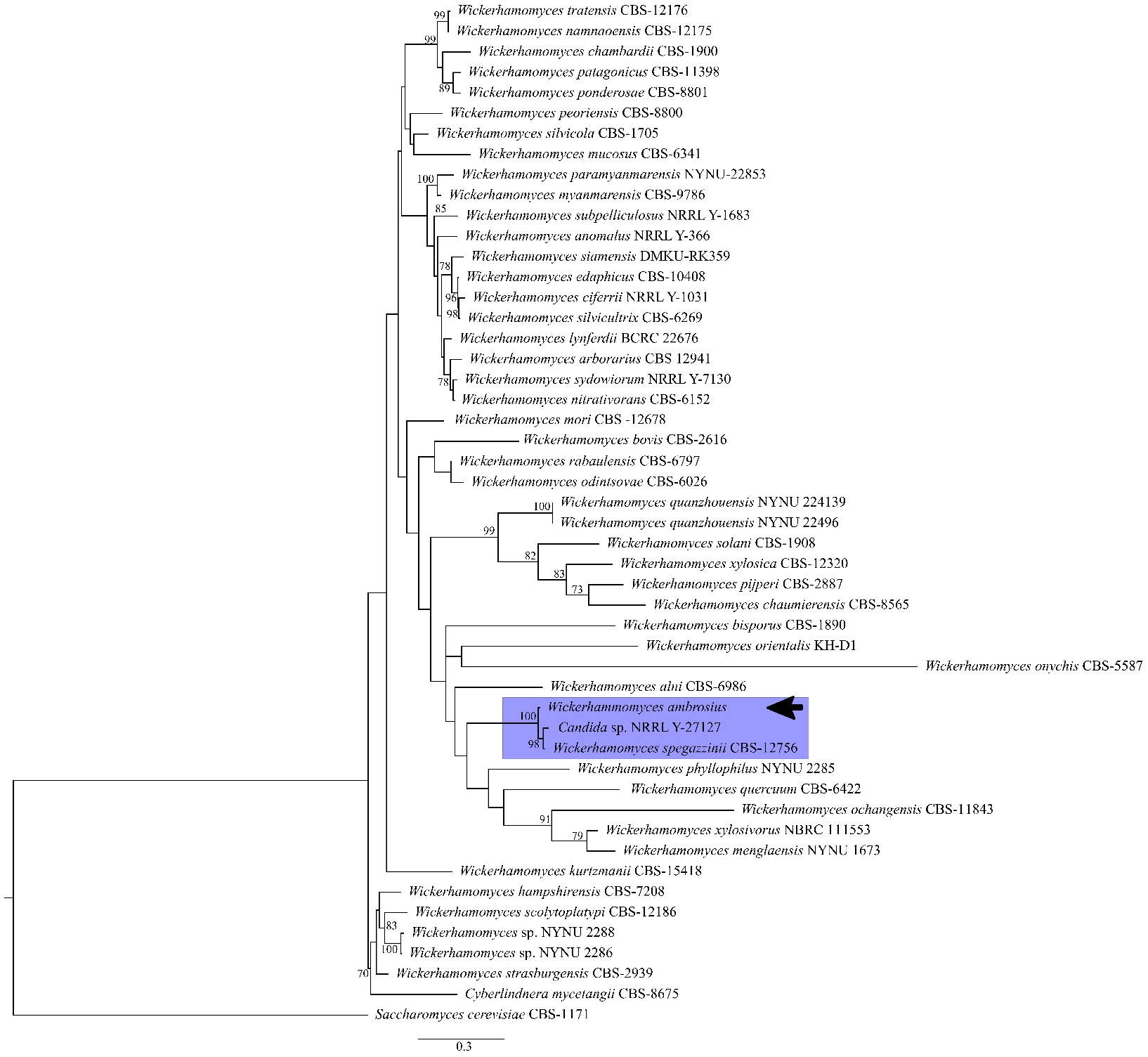

**Fig. S21**: Phylogenetic placement of *Wickerhamomyces ambrosius*. A maximum-likelihood phylogenetic tree was inferred from *ITS* gene sequences. Numbers at nodes indicate bootstrap support values (RaxML). *Schizosaccharomyces pombe* was used as the outgroup. The scale bar represents substitutions per site.

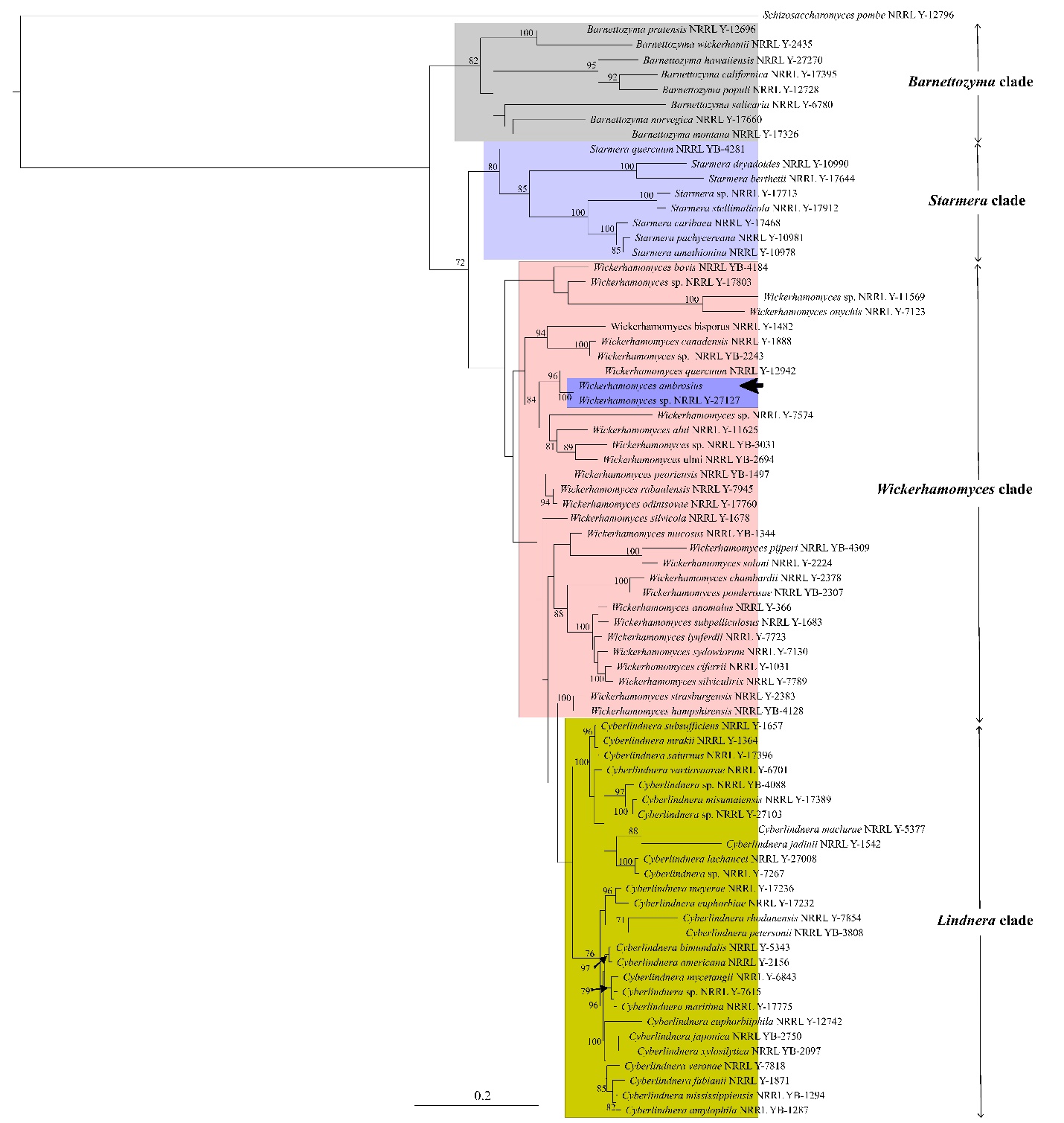

**Fig. S22**: Phylogenetic placement of *Wickerhamomyces ambrosius*. A maximum-likelihood phylogenetic tree was inferred from *LSU* gene sequences. Numbers at nodes indicate bootstrap support values (RaxML). *Schizosaccharomyces pombe* was used as the outgroup. The scale bar represents substitutions per site.

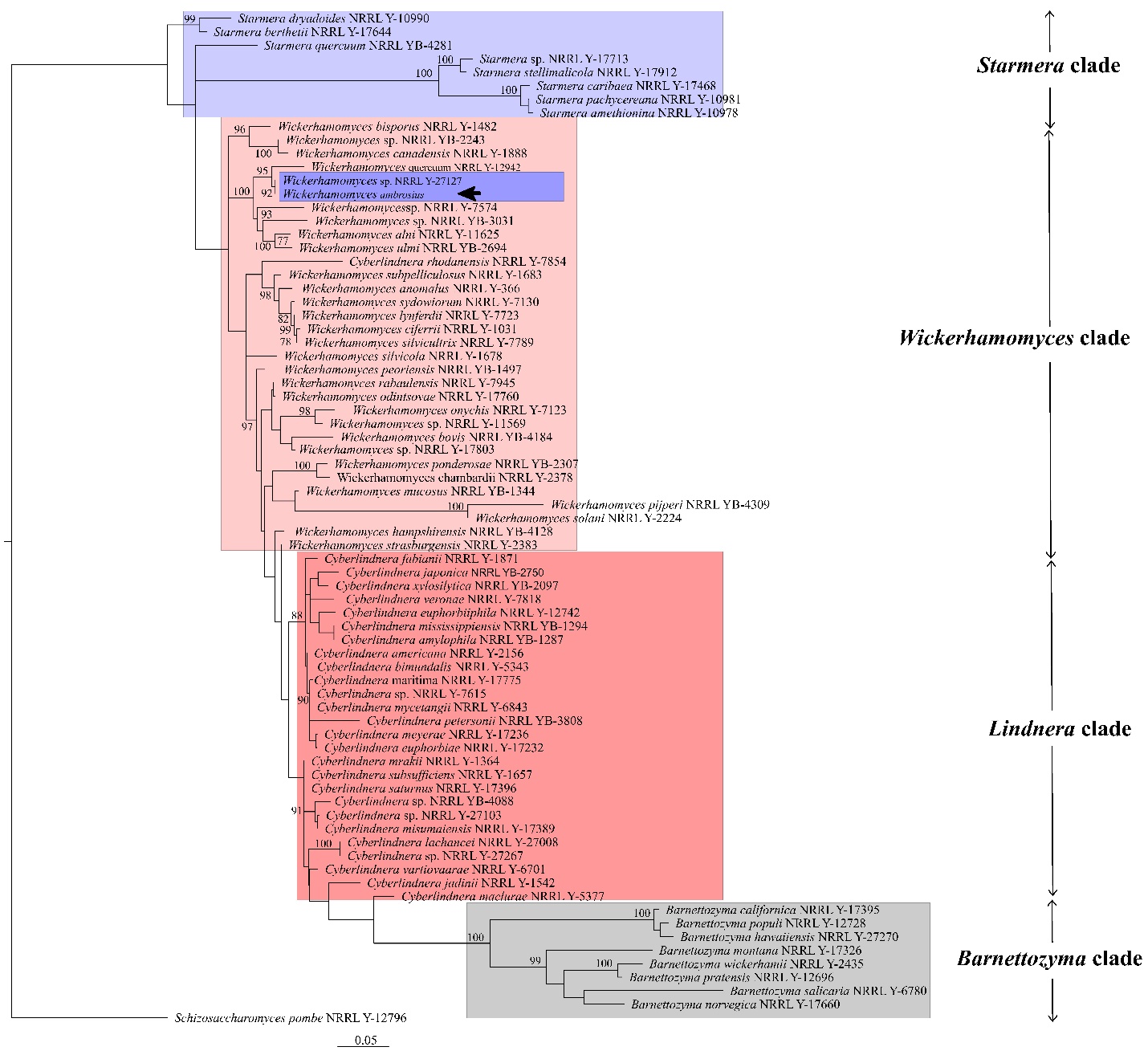

**Fig. S23**: Phylogenetic placement of *Wickerhamomyces ambrosius*. A maximum-likelihood phylogenetic tree was inferred from *SSU* gene sequences. Numbers at nodes indicate bootstrap support values (RaxML). *Schizosaccharomyces pombe* was used as the outgroup. The scale bar represents substitutions per site.

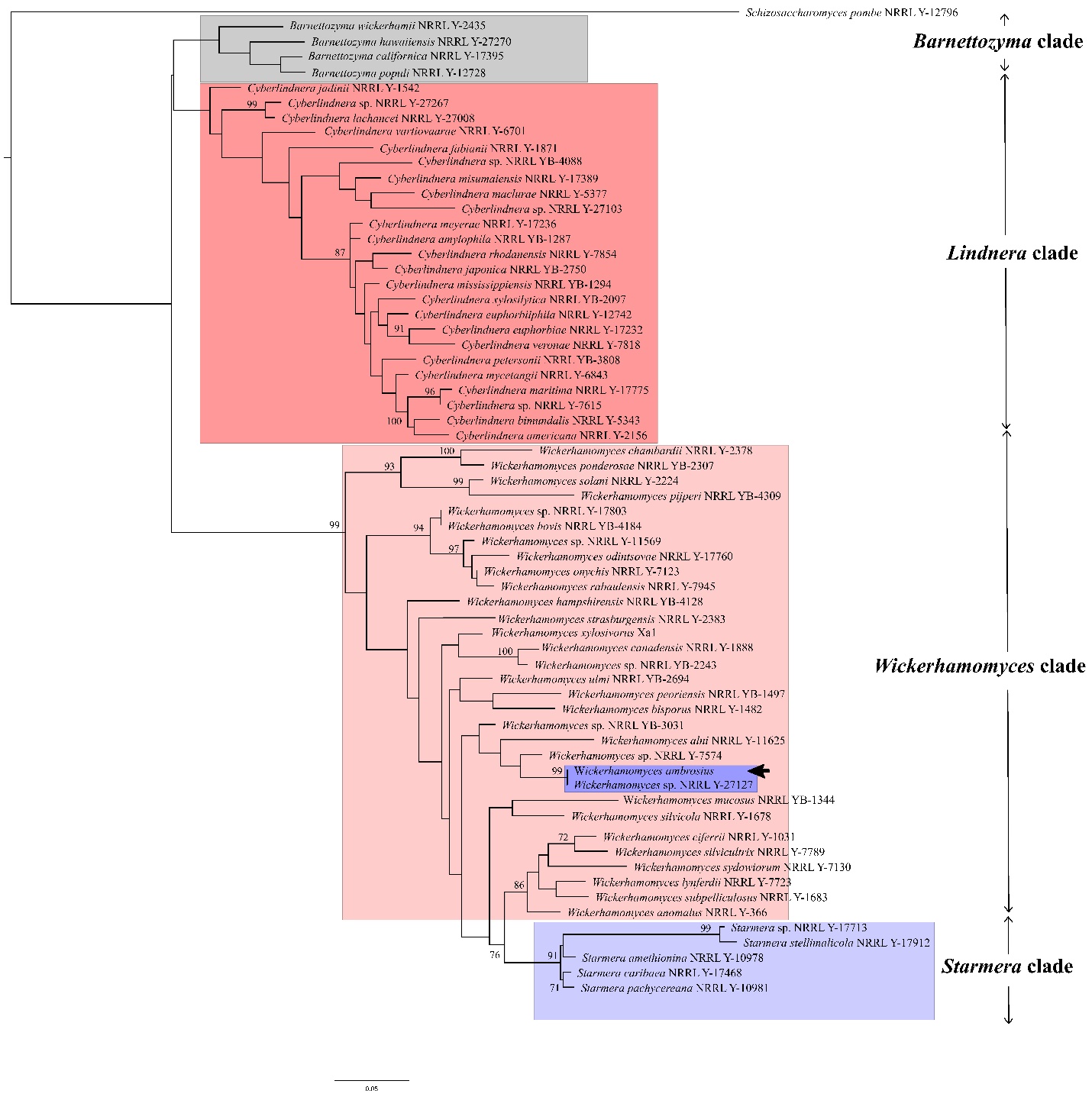

**Fig. S24**: Phylogenetic placement of *Wickerhamomyces ambrosius*. A maximum-likelihood phylogenetic tree was inferred from *TEF* gene sequences. Numbers at nodes indicate bootstrap support values (RaxML). *Schizosaccharomyces pombe* was used as the outgroup. The scale bar represents substitutions per site.

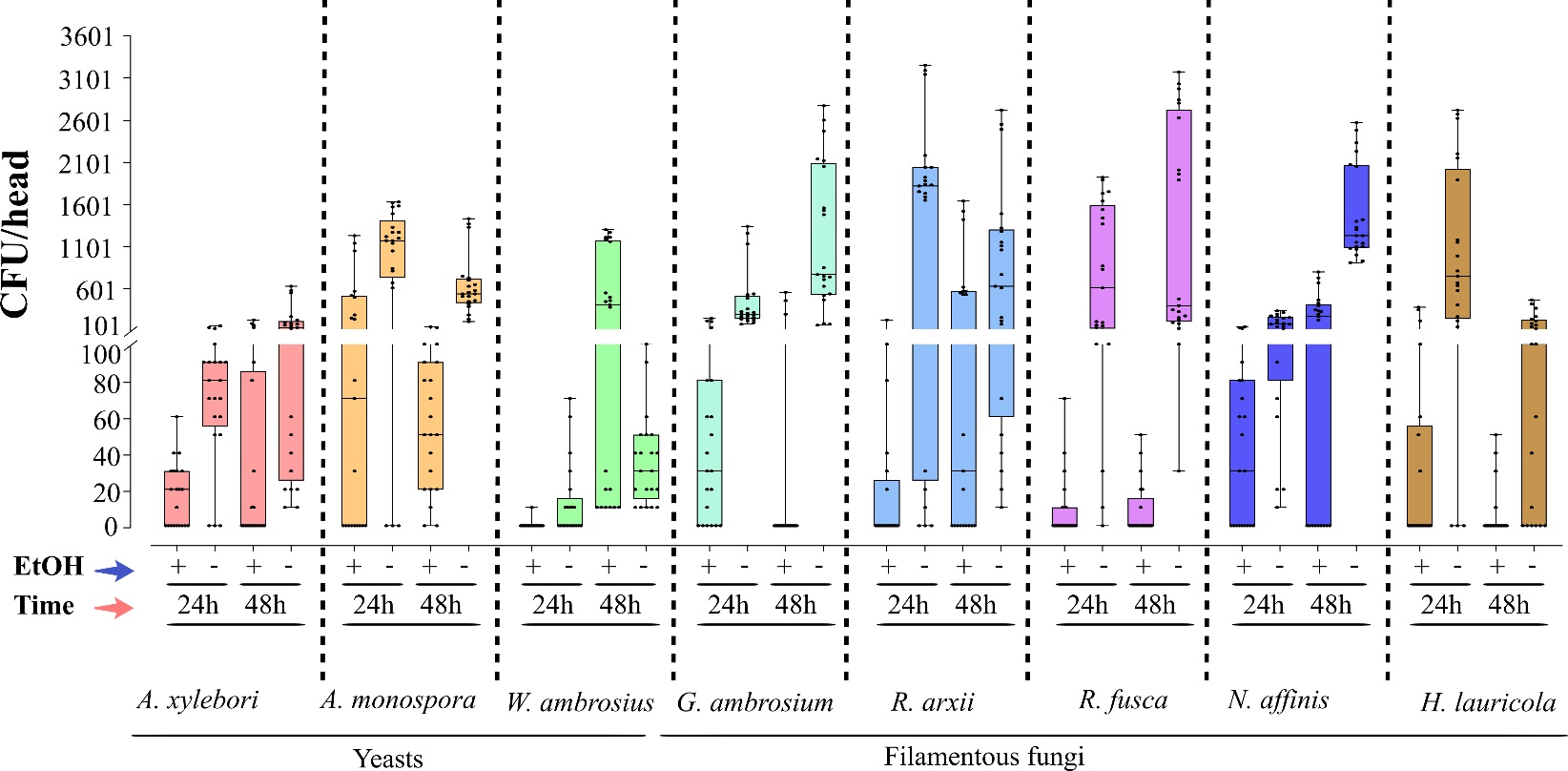

**Fig. S25**: Mycangial colonization efficiency of yeast and filamentous fungal associates in aposymbiotic ambrosia beetles. Mycangial colonization bioassays were conducted by individually exposing aposymbiotic adult beetles to PDA wells pre-inoculated with test fungi for 24 or 48 h. Following exposure, beetle heads were removed, surface-treated with or without ethanol (EtOH), homogenized, and plated to quantify fungal recovery as detailed in the Methods section. Seven fungal isolates were evaluated, including yeast-like fungi (*Alloascoidea xylebori*, *Ambrosiozyma monospora*, *Wickerhamomyces ambrosius*) and filamentous fungi (*Graphium ambrosium*, *Raffaelea arxii,* *Raffaelea fusca*, and *Neocosmospora affinis*). *Harringtonia lauricola* was included as a positive control due to its established ability to colonize beetle mycangia. *Candida albicans* was used as a negative control. Data for *C. albicans* showed <5 CFUs/head and is omitted from the graph. Boxplots represent the distribution of raw data (CFU/head) values, with individual data points shown.

| **Table Supplemental S1**: Dataset used for phylogenomic analyses in this study. Sequences for the indicated loci were derived from the given references. | | |
| --- | --- | --- |
| **Taxa** | **Loci** | **References** |
| *Neocosmospora affinis* | *ITS*, *LSU*, *RPB1*, *RPB2* and *TEF* | Carrillo JD et al., 2019  Lynn KMT et al., 2021  Sandoval-Denis M et al., 2019  Zhang YX et al., 2024 |
| *Graphium ambrosium* | *SSU*, *β-tubulin* (*BTUB*), *LSU*, *ITS*, and *TEF* | Bangash NK et al., 2025  Paciura D et al., 2010 |
| *Alloascoidea xylebori* | *LSU*, *SSU*, *TEF*, *RPB1*, and *RPB2* | Kurtzman CP et al., 2013a  Kurtzman CP et al., 2013b |
| *Wickerhamomyces ambrosius* | *ITS*, *LSU*, *SSU*, and *TEF* | Kurtzman CP et al., 2008  Chai CY et al., 2014 |
| *Raffaelea arxii* | *LSU* | Simmons DR et al., 2016  Dreaden TJ et al., 2014 |
| *Raffaelea fusca* | *LSU* | Simmons DR et al., 2016  Dreaden TJ et al., 2014 |
| *Ambrosiozyma monospora* | *TEF* | Kurtzman CP et al., 2013a  Kurtzman CP et al., 2013b |

| **Supplemental Table S2:** The list of primers and PCR targets used in this study | | | |
| --- | --- | --- | --- |
| **Target loci** | **Primer name** | **Primer sequence (5′-3′)** | **References** |
| *ITS* rDNA | ITS_F | GCATGTGTTGATTATGTCCC | Designed in this study |
|  | ITS_R | AAGCATCTTTTCCTCCGCTT |  |
| β-tubulin (*BTUB*) | T1 | AACATGCGTGAGATTGTAAGT | Kerry O'Donnell, Elizabeth Cigelnik (1997). |
|  | T22 | TCTGGATGTTGTTGGGAATCC |  |
| 3′-*TEF1*-α | 983F | GCYCCYGGHCAYCGTGAYTTYAT | Rehner, Stephen A., and Ellen Buckley (2005) |
|  | 2218R | ATGACACCRACRGCRACRGTYTG |  |
| *LSU* (D1/D2) | NL-1F | GCATATCAATAAGCGGAGGAAAAG | Kurtzman, C. P., & Robnett, C. J. (2013) |
|  | NL-4R | GGTCCGTGTTTCAAGACGG |  |
| *SSU* rDNA | NS-1AF | GCCAGTAGTCATATGCTTGTCTC | White, T.J. (1990) |
|  | NS-8R | TCCGCAGGTTCACCTACGGA |  |
| *RPB1* | DF2asc | CAYAAGGARTCYATGATGGGTCAT | [Kerry O’Donnell](https://www.sciencedirect.com/author/7101682936/kerry-l-o-donnell) et al. (2013) |
|  | G2R | GTCATYTGDGTDGCDGGYTCDCC |  |
| *RPB2* | 5F | GAYGAYMGWGATCAYTTYGG | [Kerry O’Donnell](https://www.sciencedirect.com/author/7101682936/kerry-l-o-donnell) et al. (2013) |
|  | 7cR | CCCATRGCTTGYTTRCCCAT |  |
| *RPB2* (exons 4-9) | YRPB2-4F | GGCYACTGGTAAYTGGGGTG | Kurtzman, C. P., & Robnett, C. J. (2013) |
|  | YRPB2-9R | GCAAATTTRTCACCAATTTGKGG |  |
| *RPB2* (exons 6-11) | YRPB2-6F | CWGATGCWGGTRGWGTTTAYMGWC | Kurtzman, C. P., & Robnett, C. J. (2013) |
|  | YRPB2-11R | CTTATCATCCACCATATGTC |  |
| Note: Overlapping *RPB2* fragments from yeasts were assembled to generate the full *RPB2* sequence. | | | |

| **Supplemental Table S3:** PCR cycle conditions for primers used in this study^1^ | | | | | |
| --- | --- | --- | --- | --- | --- |
| **Target locus** | **Expected size (bp)** | **Initial denaturation** | **Cycling conditions** | **Final extension** | **PCR type** |
| ***ITS* rDNA** | ~1,000 | 94 °C, 1 min | 35 × (94 °C 30 s, 48 °C 30 s, 68 °C 2 min) | 68 °C, 10 min | Standard |
| **β-tubulin (*BTUB*)** | ~1,850 | 94 °C, 2 min | 35 × (94 °C 35 s, 52 °C 55 s, 72 °C 1 min) | 72 °C, 10 min | Standard |
| **3′-*TEF1*-α** | ~1,200 | 94 °C, 2 min | 10 × touchdown (66→56 °C), then 40 × (94 °C 30 s, 56 °C 30 s, 72 °C 1 min) | 72 °C, 10 min | Touchdown |
| ***LSU* (D1/D2)** | 550-600 | 94 °C, 3 min | 35 × (94 °C 1 min, 60 °C 1 min, 72 °C 1 min) | 72 °C, 10 min | Standard |
| ***SSU* rDNA** | ~1,700 | 94 °C, 5 min | 35 × (94 °C 1 min, 52 °C 1 min, 72 °C 1 min) | 72 °C, 10 min | Standard |
| ***RPB1*** | ~1,000-1,200 | 94 °C, 3 min | 35 × (94 °C 1 min, 52 °C 1 min, 72 °C 1 min) | 72 °C, 10 min | Standard |
| ***RPB2*** | ~900-1,200 | 94 °C, 3 min | 35 × (94 °C 1 min, 52 °C 1 min, 72 °C 1 min) | 72 °C, 10 min | Standard |

^1^ All PCR reactions were conducted in a total volume of 25 μL, containing 6.5 μL of 2× Taq PCR SuperMix (APExBIO, Houston, TX, USA), 2.5 μL of each primer, 2 μL of template DNA, and sterile distilled water to bring the volume to 25 μL. Amplifications were performed on a C1000 Touch thermal cycler (Bio-Rad Laboratories, Hercules, CA, USA). PCR products were visualized on 1% agarose gels, and amplicons yielding single bands of the expected size were purified using the GeneJET PCR Purification Kit (Thermo Fisher Scientific, Waltham, MA, USA).

| **Supplemental Table S4**: GenBank accession numbers for genomic loci produced in this study | | | | | | | | |
| --- | --- | --- | --- | --- | --- | --- | --- | --- |
| **Genes** | *A. xylebori*  NRRL 65231^1^ | | *A. monospora*  NRRL 65235 | *R. arxii*  NRRL 65233 | *G. ambrosium*  NRRL 65230 | *N. affinis*  NRRL 65229 | *R. fusca*  NRRL 65234 | *W. ambrosius*  NRRL 65232 |
| ***18S*** | | PX625444 | PX699942 | PX630596 | PX630594 | PX630597 | PX630595 | PX630598 |
| ***28S*** | | PX630604 | PX630682 | PX630607 | PX630606 | PX630605 | PX630608 | PX630609 |
| ***RPB1*** | | PX719819 | NA | NA | NA | PX905775 | NA | NA |
| ***RPB2*** | | PX737453 | NA | NA | NA | PX905776 | NA | NA |
| ***TEF*** | | PX696034 | PV851425 | PX757121 | PX933391 | PV220985 | PX757122 | PX929709 |
| ***ITS*** | | PX925773 | PX925772 | PX630601 | PX630600 | PX630599 | PX630602 | PX630603 |
| ***BTUB*** | | NA | NA | PX781868 | PX868310 | NA | PX848778 | NA |
| ^1^ Cultures of all isolates have been deposited into the NRRL = Agricultural Research Service Culture Collection (NRRL), U.S. Department of Agriculture, Peoria, Illinois, USA | | | | | | | | |
